# Off-target interaction of the amyloid PET imaging tracer PiB with acetylcholinesterase

**DOI:** 10.1101/2025.05.16.654428

**Authors:** Alberto Granzotto, Rosa Fullone, Ludovico Miccoli, Manuela Bomba, Claudia Di Marzio, Stefano Delli Pizzi, Giuseppe Floresta, Stefano L. Sensi

## Abstract

Pittsburgh compound B (PiB) is a widely used Positron Emission Tomography (PET) tracer for detecting amyloid-β (Aβ) deposits in Alzheimer’s disease (AD). While PiB is assumed to bind selectively to Aβ, emerging evidence suggests off-target interactions that may complicate PET signal interpretation. Here, we report that PiB can interact with acetylcholinesterase (AchE), a key enzyme in the cholinergic system. Similarity screening identified the AchE ligand thioflavin T (ThT) as the top structural analog of PiB. Docking studies and molecular dynamics simulations showed that PiB stably binds the peripheral anionic site (PAS) of AchE, with binding energies comparable to ThT and clinically relevant AchE inhibitors. *In vitro* fluorescence-based assays confirmed this interaction and suggest an involvement of the PAS. These findings indicate a stable off-target interaction between PiB and AChE with implications for interpreting PiB-PET signals in AD, particularly in regions with altered AchE expression or under AchE inhibitor therapy.

---

Positron emission tomography (PET)-based biomarkers are largely employed in research and clinical settings for disease diagnosis and monitoring, patient stratification, or as an efficacy outcome of interventions ^1^. In Alzheimer’s disease (AD), PET tracers have been developed to quantitatively detect changes in the accumulation of key pathological markers, like cortical amyloid-β (Aβ) deposits, hyperphosphorylated tau (p-tau) protein buildup, and neurodegeneration ^2,3^. Alterations in these biomarkers mirror disease progression and are the “gold standard” for diagnosing AD and for the early detection of people at-risk of developing the condition ^4,5^. The shift from a clinical- to a biomarker-based definition of AD is also at the basis of the “ATN research framework”, a biological definition of the disease (i.e., ’A’ – amyloid, ‘T′ – tau, and ‘N′ – neurodegeneration) aimed at offering a quantifiable and unbiased staging of AD ^4^. The approach is relevant since the pathological alterations of AD can occur and are detectable long before the onset of cognitive and behavioral symptoms ^6,7^. Early identification of individuals in the very early stages of the condition represents a transformative step for the effective development and targeted implementation of disease-modifying interventions.

Alterations of Aβ levels are widely recognized as one of the earliest molecular changes that can foreshadow the onset of AD pathology, although the specific contribution of Aβ to disease pathogenesis is debated ^8,9^. Quantitative assessment of Aβ is either performed in biological fluids like liquor and plasma, where decreases in Aβ abundance reflect the cerebral deposition of the peptide, or by PET-based imaging, where specific radioligands are employed to detect the presence of fibrillar Aβ (fAβ) aggregates in the brain. Several Aβ radiotracers have been developed since the early 2000s with Pittsburgh compound B (^11^C-PiB), a thioflavin T (ThT) analog, being the first of this class of imaging agents. The short half-life of ^11^C-PiB led to the development of fluorine-18 derivatives more suitable for clinical applications, like ^18^F-flutemetamol or the trans-stilbene-based compounds ^18^F-florbetapir and ^18^F-florbetaben. Nevertheless, ^11^C-PiB is still broadly adopted in clinical research settings.

Although these radioligands are widely employed for the diagnosis of AD and for monitoring target engagement of Aβ-targeting interventions, doubts have been cast on their specificity and sensitivity ^10–13^. Previous studies demonstrated that 2-aryl-6-hydroxybenzothiazole-based tracers can effectively bind to off-target molecules, like sulfotransferases, that likely contribute to PET signals unrelated to the overall Aβ load ^10,12^. However, it is unclear whether this class of Aβ radioligands has additional off-target effects.

This study aims to investigate PiB binding characteristics at the molecular level and, by employing unbiased *in silico* screening, docking calculations, molecular dynamics (MD) simulations, and *in vitro* assay, we surveyed for potential novel binding partners unrelated to Aβ pathology.

To identify potential off-target partners of Aβ PET tracers, we performed an unbiased screening of biologically relevant molecules that show structural similarity with PiB by employing the SwissSimilarity 2021 Web Tool. Our analysis returned the score of 400 molecules (Fig. 1A and Supplementary chart 1) with ThT being the top-scoring molecule (score 0.996). More importantly, ThT was identified because the molecule is a ligand for acetylcholinesterase (AchE) in the Protein Data Bank (PDB ID: pdb_00002j3q). To test the hypothesis that PiB interacts with AchE, we performed docking studies within the AchE pocket using the crystal structure of the human AchE (PDB ID: pdb_00004ey7). AchE has two binding sites: the catalytic site and the peripheral anionic site (PAS) ^14,15^. We focused on the latter, located at the entrance of the catalytic gorge, since it mediates the interaction of AchE with ThT ^16–18^. Analysis of docking results shows that PiB has binding energy comparable with that of ThT (8.603 and 8.771 kcal/mol, respectively; Fig. 1B). Binding energy of clinically approved AchE inhibitors (donepezil, galantamine, and rivastigmine) was calculated for comparison (Fig. 1B). Fig. 1C and D show the two- and three-dimensional poses and the interaction of PiB with the amino acid residues in the AchE PAS. PiB forms a π–π-sulfur interaction with residue Phe297 along with several hydrophobic, alkyl, and Van der Waals interactions with residues Trp86, His447, Tyr337, Phe338, Tyr72, Trp286, Phe295, Tyr341, and Tyr124 (Fig. 1C, D).

**Figure 1.**
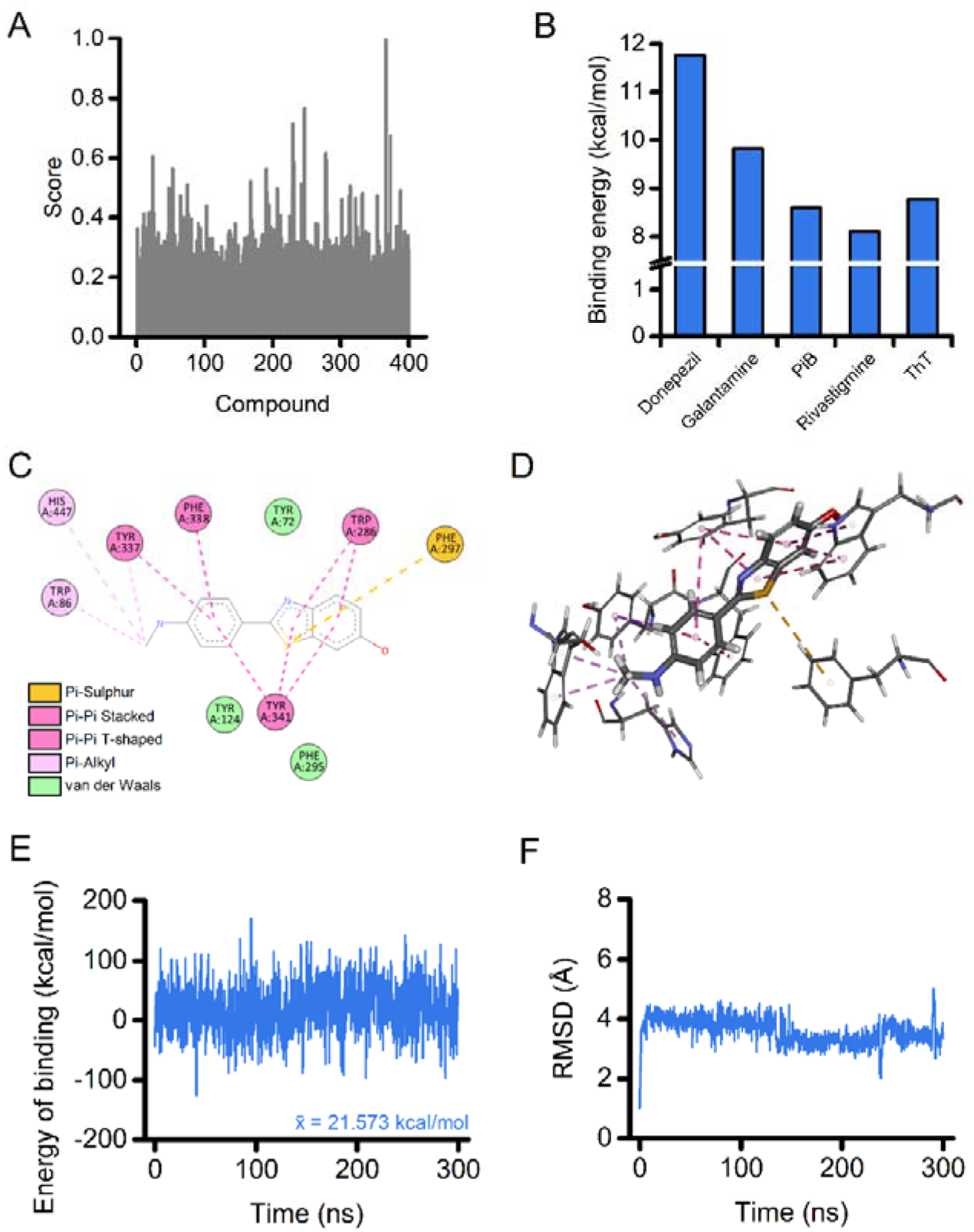
Identification and *in silico* characterization of AchE as a potential target of PiB. (A) The plot illustrates the similarity score of each compound screened with the SwissSimilarity 2021 Web Tool. (B) The histogram depicts the binding energy calculation of the listed ligands after docking on AchE. (C - D) Two-dimensional (C) and three-dimensional (D) docking poses and interactions of PiB in the AchE PAS. The dashed yellow line indicates π–π-sulfur interaction; dashed pink lines indicate π–π-alkyl interactions; magenta lines indicate π–π and T- shaped interactions; green residues show van der Waals interactions. (E - F) Time course of energy of binding (E) and root mean squared displacement (RMSD; F) for the PiB-AchE complex over a 300 ns MD simulation.

We further investigated the PiB-AchE complex by performing a 300 ns molecular dynamics (MD) simulation. Analysis of the energy of binding shows that PiB maintains a high and stable binding energy throughout the simulation (Fig. 1E). The stability of the PiB-AchE complex is also supported by the root mean square deviation (RMSD) analysis of the ligand movement after superimposing the molecule on the enzyme structure (Fig. 1F). After a stabilization phase, the ligand remains within the AchE PAS. The compound exhibits only a few modest, sharp fluctuations that return to baseline levels during the simulation (Fig. 1F). We attribute the stability of the complex to the sulfur interaction, along with the dense network of hydrophobic interactions that keep PiB within the AchE PAS.

To determine whether PiB forms a stable complex with AchE *in vitro*, we leveraged the intrinsic spectroscopic properties of the compound. The molecule displays an absorbance and an emission maximum at 348 and 432 nm, respectively (Fig. 2A). We measured the fluorescence signal of PiB after incubation with or without AchE from *Electrophorus electricus* (eeAchE), followed by size-exclusion filtration to remove unbound ligand. Samples incubated with eeAchE retained a significantly higher fluorescence signal when compared to the PiB sample alone, indicating PiB binding to the enzyme (Fig. 2B and C). The specificity of this approach was confirmed by performing a similar set of experiments in which PiB was incubated with bovine catalase, an enzyme that shares with eeAchE a tetrameric structure and a similar molecular weight (Fig. 2D and E).

**Figure 2.**
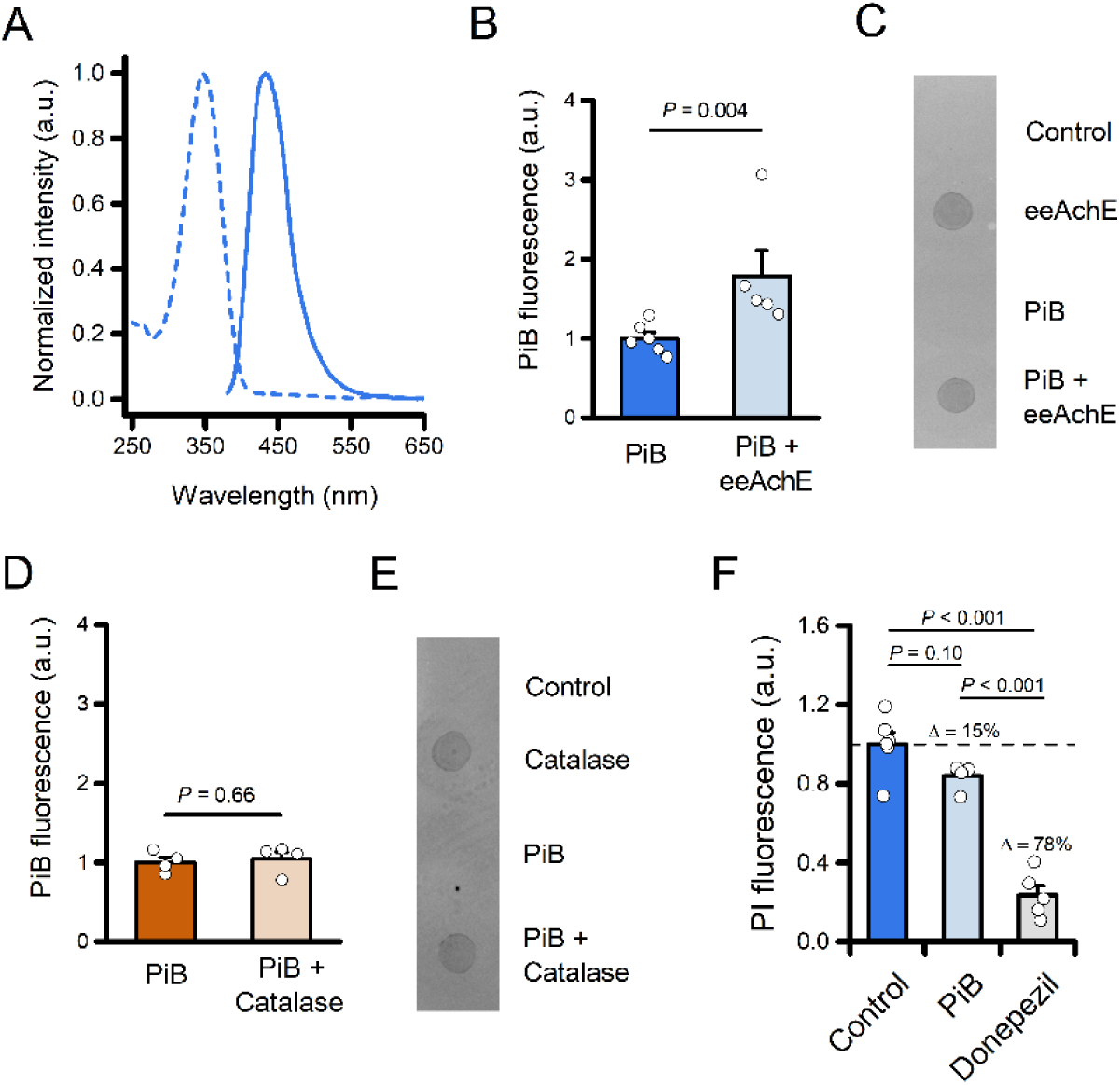
Experimental validation of the formation of the PiB-eeAchE complex. (A) Absorption (dashed line) and emission (solid line) normalized spectra of PiB in KPi buffer (pH 7.4). (B) The bar graph depicts normalized fluorescence of PiB following incubation of the compound with or without eeAchE (2 µg) and size-exclusion filtration (PiB n = 6 and PiB + eeAchE n = 5 independent experiments). (C) Ponceau S staining of the retentate was spotted onto a nitrocellulose membrane to assess protein recovery. (D) The bar graph depicts normalized fluorescence of PiB following incubation of the compound with or without Catalase (2 µg) and size-exclusion filtration (PiB n = 4 and PiB + Catalase n = 4 independent experiments). (E) Ponceau S staining of the retentate was spotted onto a nitrocellulose membrane to assess protein recovery. (F) The bar graph depicts normalized fluorescence of PI following incubation with eeAchE in the presence of vehicle (Control; 0.8% DMSO), PiB (20 µM, 0.8% DMSO), or Donepezil (20 µM, 0.8% DMSO). Note the ≈ 15% signal reduction in the presence of PiB. In B and D, the comparison of mean values was assessed by the Mann-Whitney U Test. In F, mean values were compared by one-way ANOVA followed by Tukey’s post-hoc test.

To further evaluate PiB binding to the PAS of eeAChE, we performed propidium iodide (PI) displacement, an assay used for probing the interaction of candidate drugs with the PAS of AchE^19^. The binding of PI to the PAS increases dye fluorescence. Meanwhile its displacement by PAS-interacting compounds leads to signal reduction ^20^. In the presence of 20 µM PiB, PI fluorescence was reduced by approximately 15% compared to control conditions (Fig. 2F). Although this difference did not reach statistical significance (*P* = 0.10), the data suggest a potential trend toward PI displacement. The reduction increased to ≈ 78% in the presence of the high-affinity, PAS-binding AchE inhibitor donepezil (20 µM; Fig. 2F) ^14,21^.

Together, these findings support the formation of a stable PiB–eeAchE complex *in vitro* and are consistent with the possibility that PiB interacts with the PAS of the enzyme.

In this study, we provide computational and experimental evidence that the amyloid PET tracer PiB can bind AchE, suggesting a previously unrecognized off-target interaction. Our findings extend prior observations that ThT-based compounds may interact with non-amyloid targets and raise important questions on the specificity of PiB and related PET tracers used in AD research and diagnostics ^16^.

Our *in silico* similarity screening identified ThT – a known AchE ligand – as the compound most structurally related to PiB among biologically relevant molecules in the PDB, pointing to a potential interaction between PiB and AchE. We further tested this hypothesis using molecular docking and MD simulations. Docking results showed that PiB binds the PAS of AchE with binding energy comparable to ThT and within the range of clinically relevant AchE inhibitors. PiB establishes π–sulfur and hydrophobic interactions with residues located in the PAS and near the gorge of the active site, like Phe297, Trp286, Tyr337, and His447. These residues were found to be key for the interaction with AchE-targeting drugs ^14^. MD simulations further confirmed the persistence of these interactions, indicating a stable and energetically favorable complex.

We validated the computational predictions with a binding assay that exploits the intrinsic fluorescence properties of PiB, supporting the formation of a stable PiB–AchE complex *in vitro* and suggesting that the interaction occurs at the PAS of the enzyme. It is important to notice that a more thorough investigation of the interaction between PiB and AchE was hampered by both the physicochemical properties of PiB and the sensitivity of AchE to PiB-compatible solvents.

We found that in conditions suitable for the AchE enzymatic assay, PiB began to precipitate at concentrations above 25 µM (Supplementary Fig. 1B). The use of alternative solvents or surfactants was unsuccessful (unpublished observations). Moreover, higher concentrations of DMSO were shown to substantially impair AchE activity ^22^. In a further attempt to directly examine PiB-PAS interaction, we also tested an *in vitro* competition assay in the presence of donepezil ^14,21^. However, preliminary control experiments showed a substantial spectral overlap between PiB and donepezil (Supplementary Fig. 1B), making fluorescence-based comparisons difficult to interpret and prone to bias. While these technical constraints limit our ability to perform orthogonal or competitive binding assays, they do not undermine the core observation that PiB interacts with AchE, as suggested in our fluorescence-based filtration assay and PI displacement.

The identification of AChE as a potential off-target of PiB has several implications. First, it imposes the need to carefully interpret PET signals in brain regions where AchE is abundantly expressed, particularly in early-stage or atypical AD presentations, where amyloid deposition may not be the unique contributor to the tracer uptake. Second, given that AchE expression and activity can change in the aging brain and neurodegenerative conditions beyond AD ^23,24^, the off- target binding of PiB to AchE could contribute to false positives or elevated baseline signals in specific populations. Third, the PiB signal could be influenced by the use of AchE inhibitors that act by binding the PAS of the enzyme, leading to false negative results. Moreover, the high lipophilicity of PiB could also explain the elevated retention of the tracer in lipid-enriched white matter regions ^25,26^.

Our findings align with previous reports of off-target binding for other radiotracers used in AD, for whom interactions with enzymes like monoamine oxidases and sulfotransferases have been reported ^10,11,27^. In addition, our similarity virtual screening does not rule out the existence of additional PiB binding partners.

To our knowledge, this is the first study to indicate an interaction between PiB and AchE at both the computational and experimental levels. While the functional consequences of this binding remain to be explored, such interactions may alter AchE activity or affect PiB signals *in vivo*. Further studies using radiolabeled PiB and AchE inhibitors *in vivo* are warranted to confirm whether this interaction occurs under pathophysiological conditions and contributes to PET signals.

In conclusion, these results underscore the importance of integrative approaches combining computational modeling with biochemical validation to uncover and assess the biological relevance of such interactions.

## Materials and methods

### Reagents and chemicals

PiB was purchased from TargetMol. eeAchE, catalase from bovine liver and all the other chemicals were from Sigma-Aldrich.

### Library screening

Similarity screening for the PiB amyloid PET tracer was performed with the SwissSimilarity 2021 Web Tool (http://www.swisssimilarity.ch/ ^28,29^) on August 3, 2024. The search was limited to ligands present in the Protein Data Bank (LigandExpo; 19500 compounds) using a consensus 2D/3D screening using a score based on both FP2 Tanimoto coefficient and Electroshape-5D Manhattan distance ^28^. Screening scores and SMILES notations were downloaded for further analysis.

### Molecular modeling

All the molecules investigated in this study were downloaded as three-dimensional conformer .sdf files from PubChem ^30^. The Energy Minimization Experiment function, using YASARA AutoSMILES for automatic force field parameter assignment, was used to optimize the 3D structure before docking.

Docking calculations were performed using VINA default docking parameters as implemented in the YASARA suite, as previously described ^31^. Briefly, the crystal structure of the target protein was downloaded from the PDB (PDB ID: pdb_00004ey7), a cell encompassing all atoms extending 5 Å from the surface of the structure of the ligand was generated, and the crystallized ligand was removed. Global ligand docking was performed using VINA using the default parameters and further refined with VINA Local Search ^32^.

The molecular dynamics simulations of the acetylcholinesterase complexes were run with the same YASARA suite ^33^ by employing the macro *md_runfast*. A cuboid periodic simulation cell extending 20 Å from the protein surface was set and filled with water (density: 0.997 g/mL). The setup included an optimization of the hydrogen bonding network ^34^ to increase the solute stability, and a pKa prediction to fine-tune the protonation states of protein residues at pH 7.4 ^35^. NaCl ions were added at a physiological concentration of 0.9%. After steepest descent and simulated annealing minimizations to remove clashes, the simulation was run for 300 nanoseconds using the AMBER14 force field ^36^ for the solute, GAFF2 ^37^ and AM1BCC ^38^ for ligands and TIP3P for water. The cutoff was 8 Å for Van der Waals forces ^39^, no cutoff was applied to electrostatic forces (using the Particle Mesh Ewald algorithm) ^40^. The equations of motions were integrated with a multiple timestep of 2.5 fs for bonded interactions and 5.0 fs for non-bonded interactions at a temperature of 298K and a pressure of 1 atm (NPT ensemble) using algorithms described previously ^41^. MD conformations were recorded every 250 ps. The energies of binding and the MD trajectory have been calculated using the *md_analyzebindenergy* macro implemented in the YASARA suite employing the MM/PBSA method as previously described^42^. Ligand movement RMSD was calculated with the YASARA *md_analyze* function after superposing on the receptor.

### PiB spectra

A 20 µM PiB (0.8 % DMSO final concentration) solution was prepared in a 100 mM potassium phosphate buffer solution (KPi, pH 7.4). Absorbance spectrum was measured by employing a PerkinElmer Lambda 35 spectrophotometer (Range: 200 – 900 nm; slit: 2nm; resolution: 1 nm; speed: 240 nm/min). Fluorescence emission spectrum was measured with a BioTek Synergy H1 plate reader (Ex λ: 350 nm; Em range: 380 – 700 nm; resolution: 1 nm; gain: 60 a.u.).

### Turbidity assay

The turbidity assay was performed as previously described ^43^. In brief, the absorbance of increasing concentrations of PiB (1.56 µM to 1600 µM) was measured at 405 nm using a PerkinElmer SPECTRAmax 190 microplate reader. Absorbance readings from the buffer alone (KPi containing 0.8% DMSO) served as reference. The purpose of the assay was to determine the highest PiB concentration that does not result in precipitation of the compound.

### Fluorescence-based interaction assay

A fluorescence-based binding assay was performed to assess the potential interaction between PiB and eeAchE. PiB (25 µM final concentration; dissolved in 100mM KPi, 0.1 % DMSO) was incubated *in vitro* either in the presence or absence of 2 µg of eeAchE in a total volume of 50 µL. Incubations were carried out for 5 hours at 30 °C under gentle agitation (1100 rpm). Following incubation, each mixture was filtered using Amicon Ultra-0.5 centrifugal filters with a 30 kDa molecular weight cutoff (Millipore), centrifuged at 14,000 × g for 10 minutes at room temperature to separate unbound PiB from AchE-bound PiB. The retentate, containing eeAchE and any bound PiB, was recovered and transferred to a black walled 96-well plate for fluorescence measurement. Fluorescence was measured using a BioTek Synergy H1 plate reader with excitation at 350 nm and emission at 440 nm. The retentate was subsequently spotted onto a nitrocellulose membrane, stained with Ponceau S, and imaged to assess protein recovery.

### Propidium iodide displacement assay

To evaluate the interaction between PiB and the PAS of eeAchE, we employed a PI displacement assay. A total of 25 U of eeAchE were incubated overnight with PI (1 µM), either alone or in the presence of PiB (20 µM, 0.8% DMSO). A parallel experiment using donepezil (20 µM, 0.8% DMSO) served as a positive control. PI fluorescence was measured using a BioTek Synergy H1 plate reader with excitation at 535 nm and emission at 630 nm.

Background fluorescence from PI alone was subtracted from all readings. Data were then normalized as F_x_/F_vehicle_, where F_x_ represents the PI fluorescence for each condition, and F_vehicle_ is the PI fluorescence in the presence of eeAchE and 0.8% DMSO.

### Statistical analysis

Microsoft Excel (Microsoft) and OriginPro (OriginLab) were employed for statistical analysis and data plotting. Data in Fig. 2 are represented as mean ± 1 standard error of the mean (s.e.m.); data points represent individual experiments. Exact P values are reported for each relevant comparison. The number of replicates and the statistical test used are provided in the figure legends.

## Author Contributions

Conceptualization, AG; methodology, AG, RF, LM, MB, CDM, SDP, and GF; formal analysis, AG and GF; investigation, AG, RF, and GF; writing original draft, AG; supervision, AG, GF, and SLS; funding acquisition, AG. The manuscript was written through contributions of all authors. All authors have given approval to the final version of the manuscript.

## Acknowledgments

AI-assisted technology (ChatGPT 4o) has been used in the writing process to improve the readability and language of the manuscript.

## Abbreviations

Aβ: amyloid-β;
AchE: Acetylcholinesterase;
AD: Alzheimer’s disease;
DMSO: dimethyl sulfoxide;
eeAchE: *electrophorus electricus* Acetylcholinesterase
KPi: Potassium phosphate buffer
PAS: Peripheral Anionic Site;
PDB: Protein Data Bank;
PET: Positron Emission Tomography (PET);
PI: Propidium iodide;
PiB: Pittsburgh compound B;
RMSD: root mean square displacement;
ThT: thioflavin T.

## Supplementary Information

**Supplementary Figure 1.**
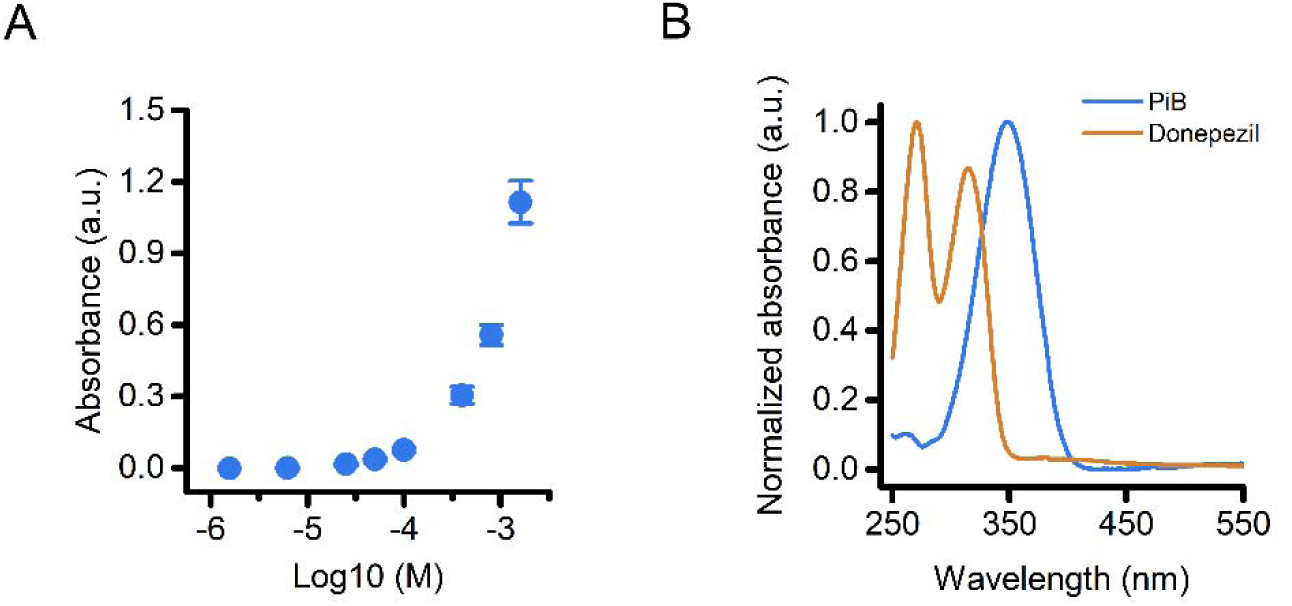
Limited PiB solubility and spectral properties of PiB and donepezil complicate fluorescence- and enzymatic-based assays. (A) The scatter plot depicts absorbance of PiB measured across increasing concentrations (1.56 – 1600 µM) and reveals precipitation above 25 µM, limiting its use in assays requiring higher concentrations. (B) Normalized absorbance spectra of PiB (blue) and donepezil (orange) highlight substantial spectral overlap which complicates interpretation of fluorescence-based competition assays involving both compounds. Data are representative of at least two independent experiments. a.u., arbitrary units.

**Supplementary chart 1.**
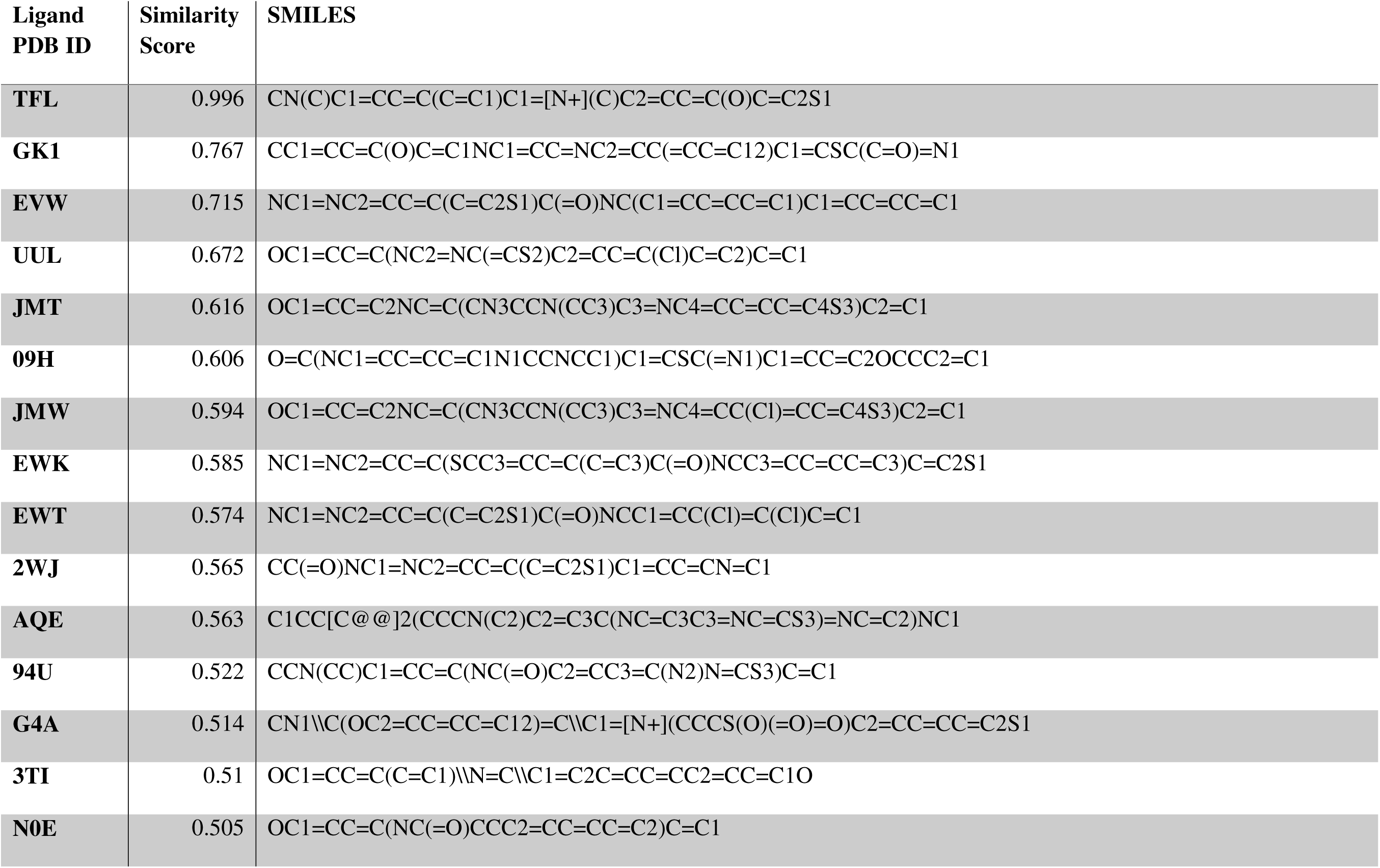
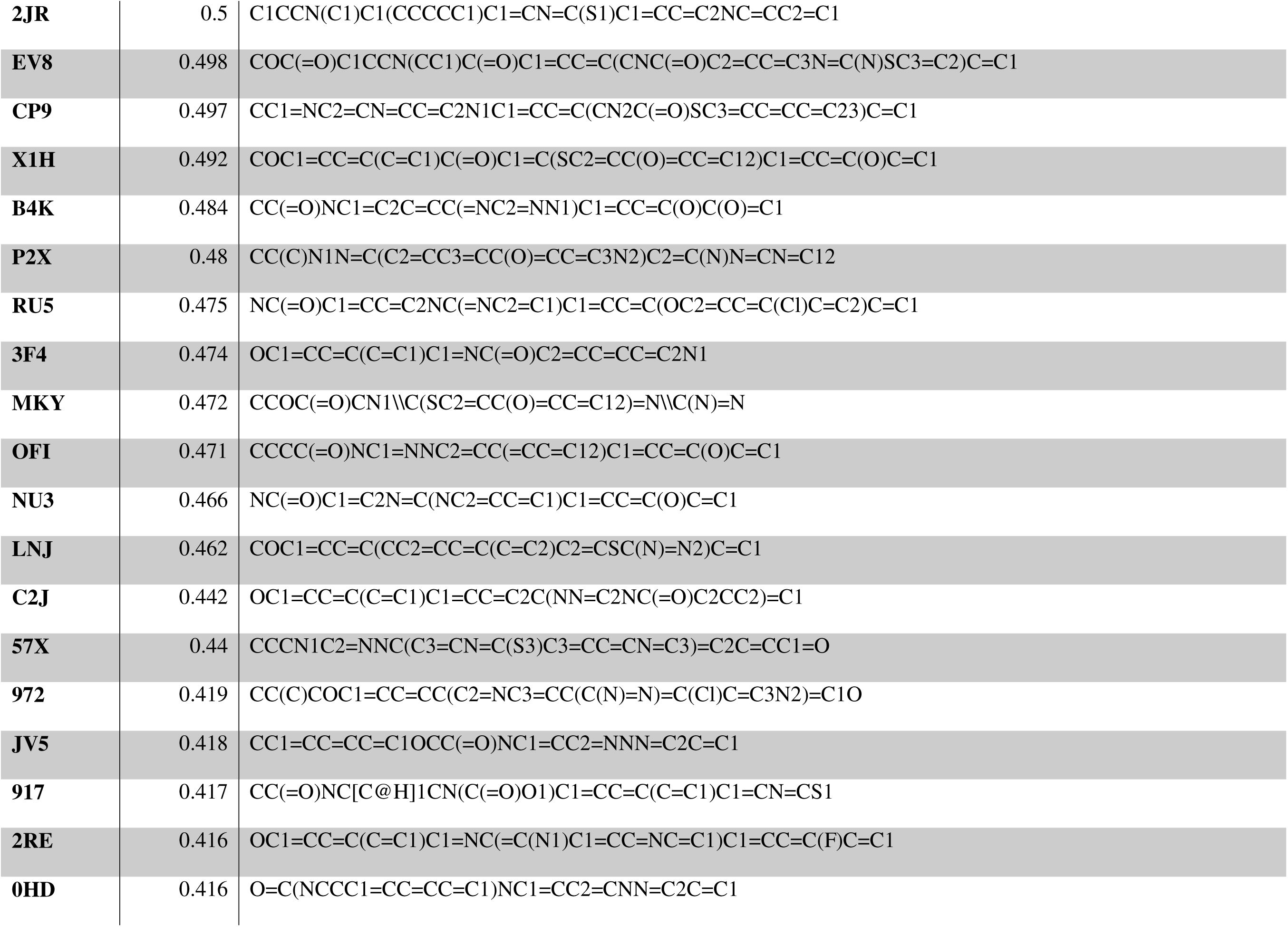
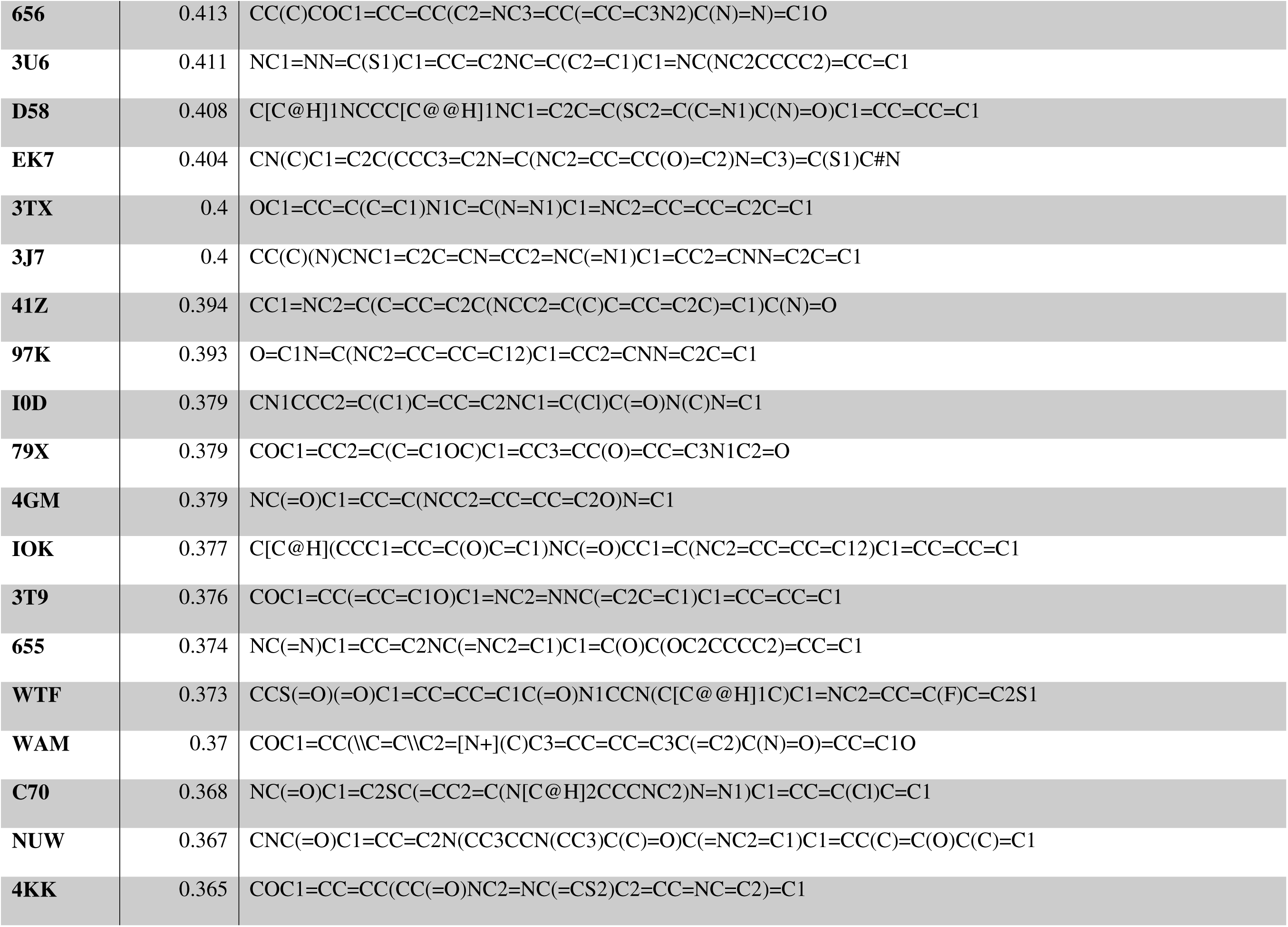
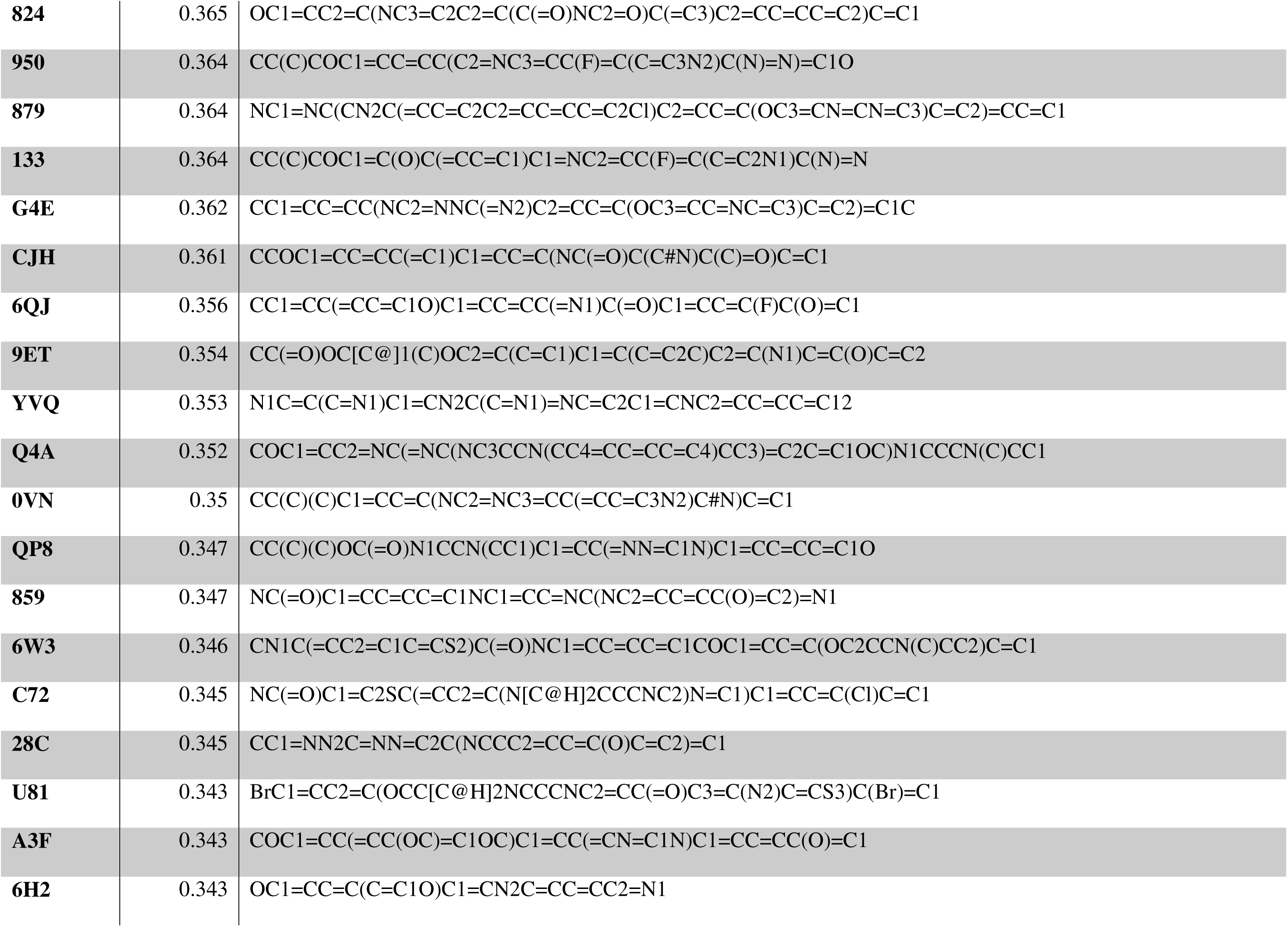
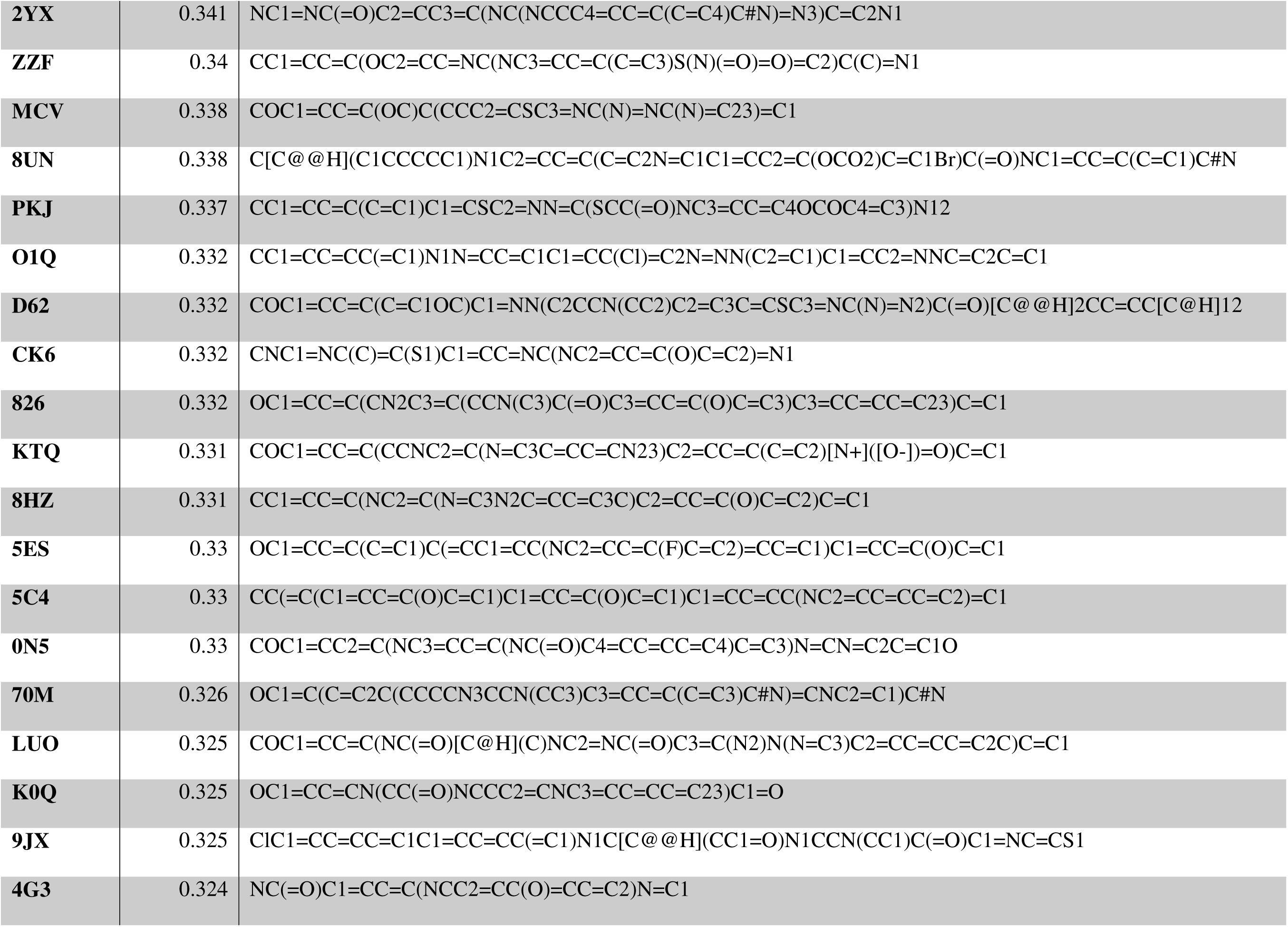
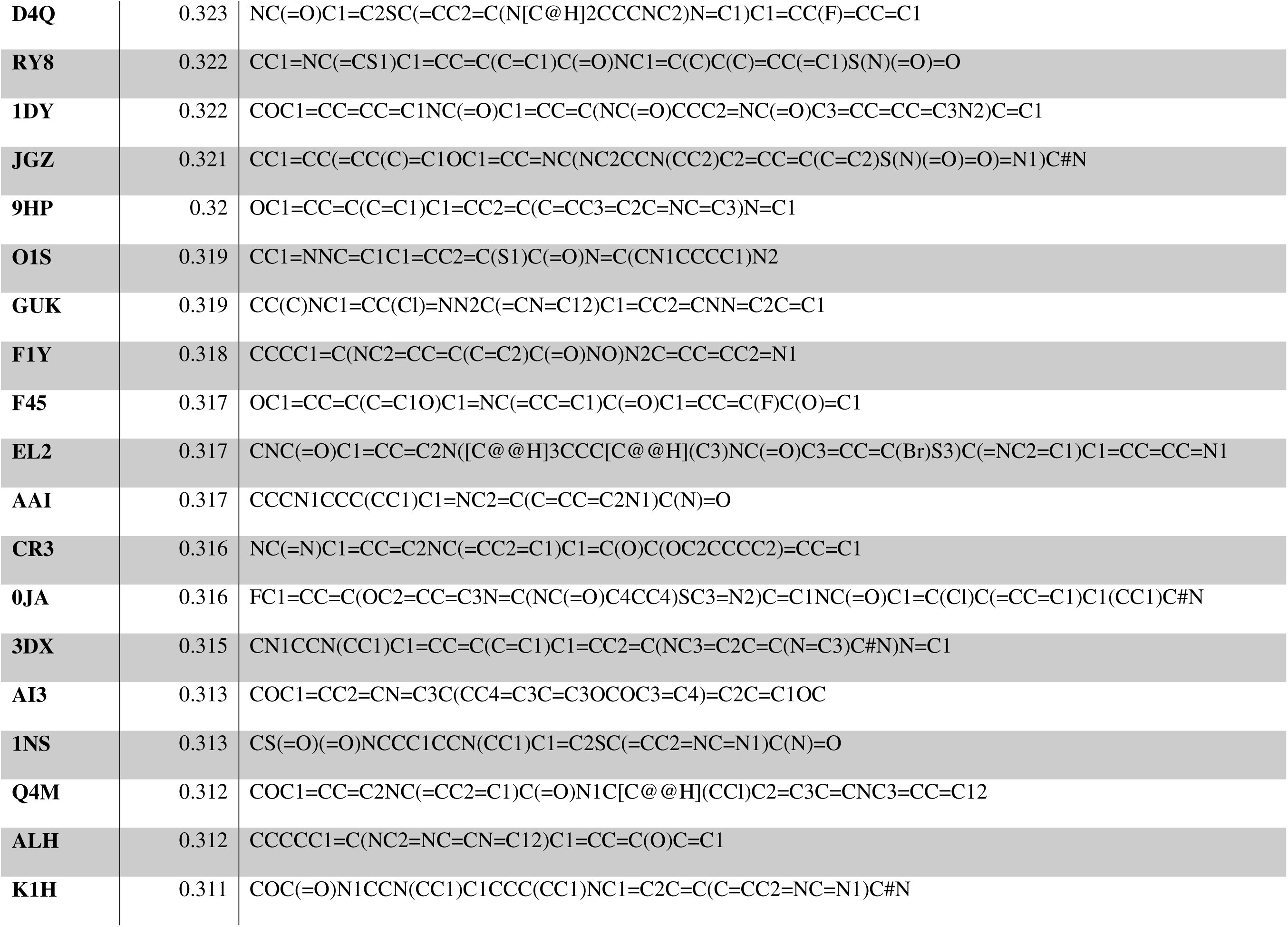
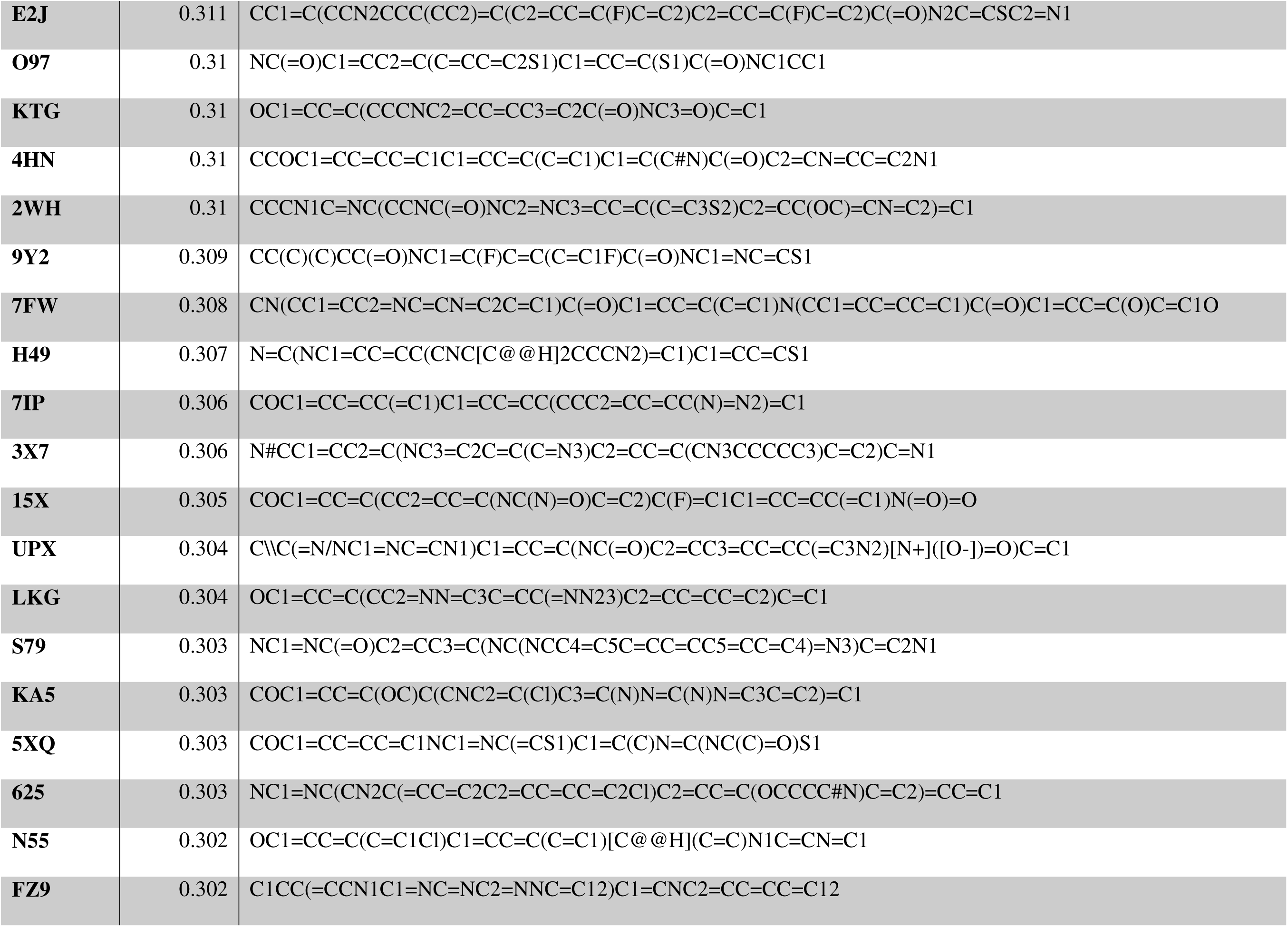
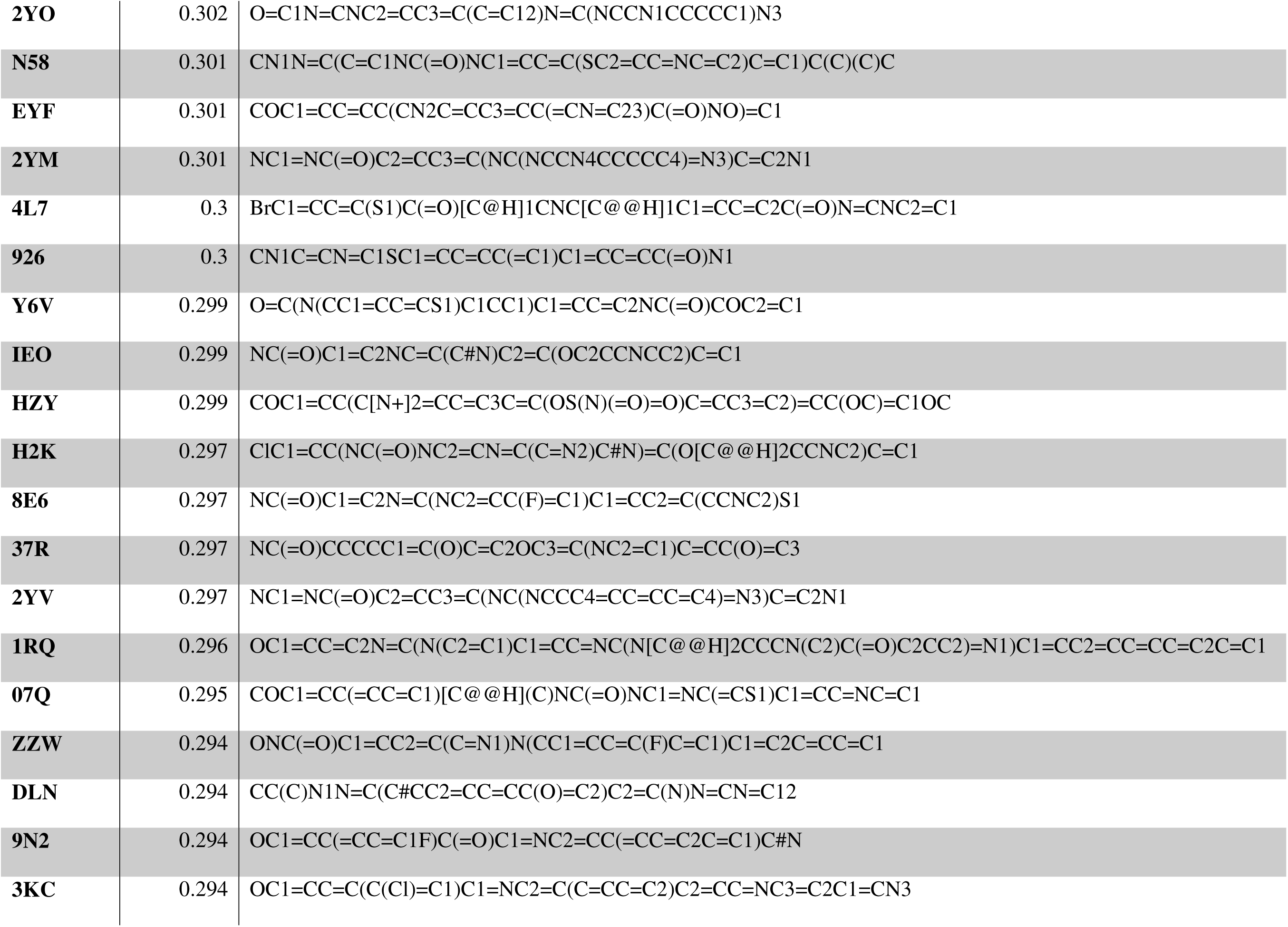
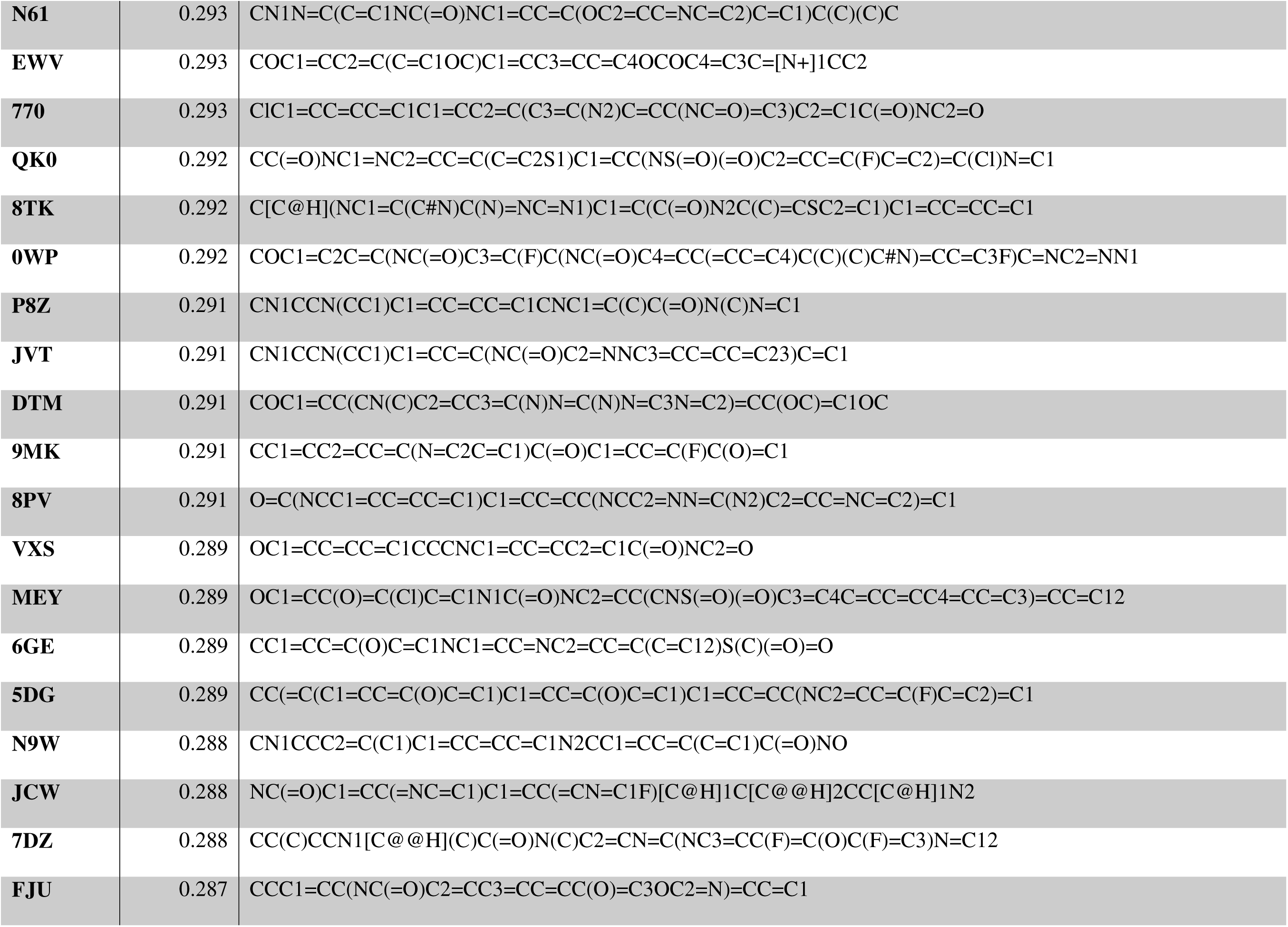
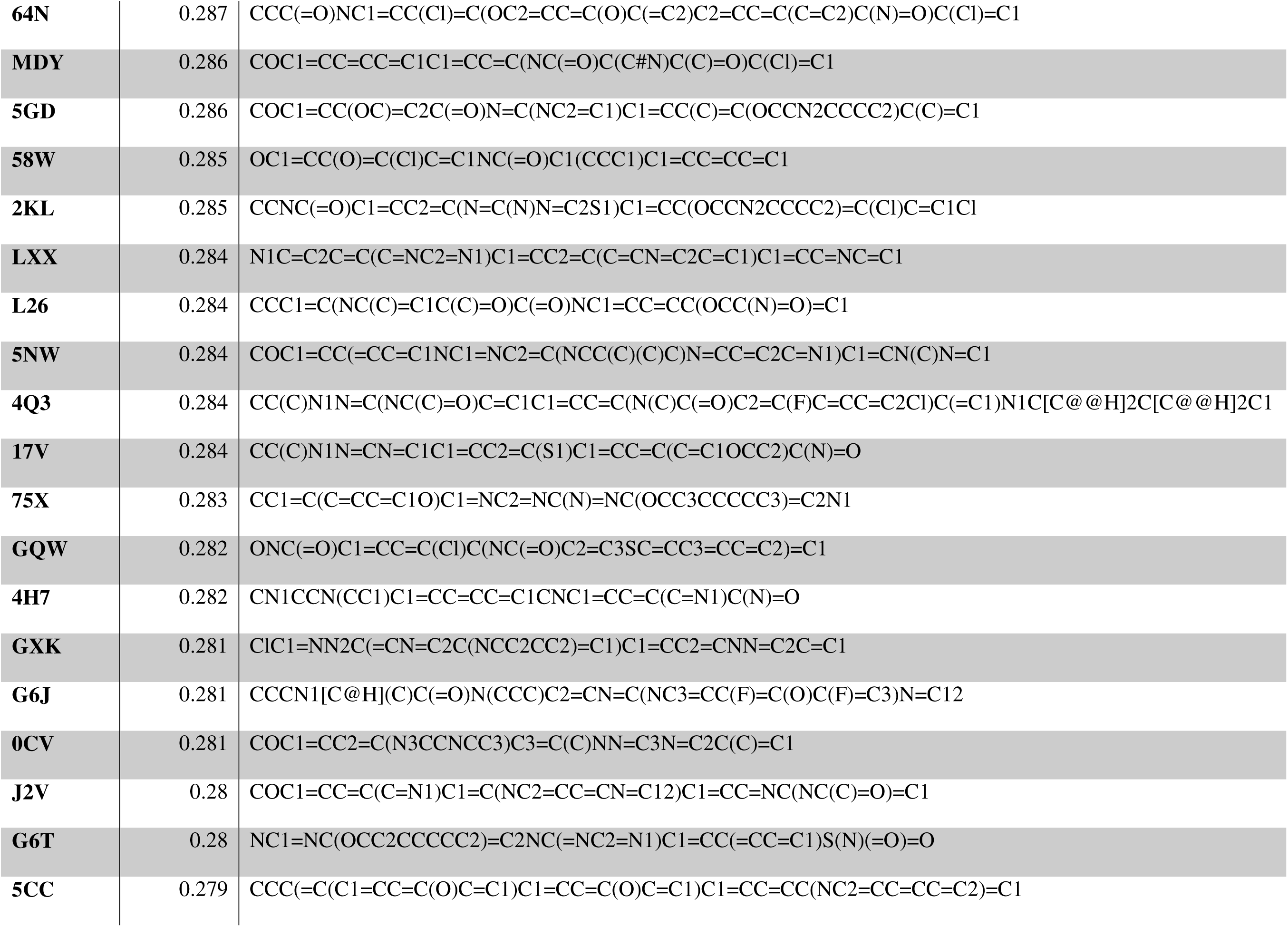
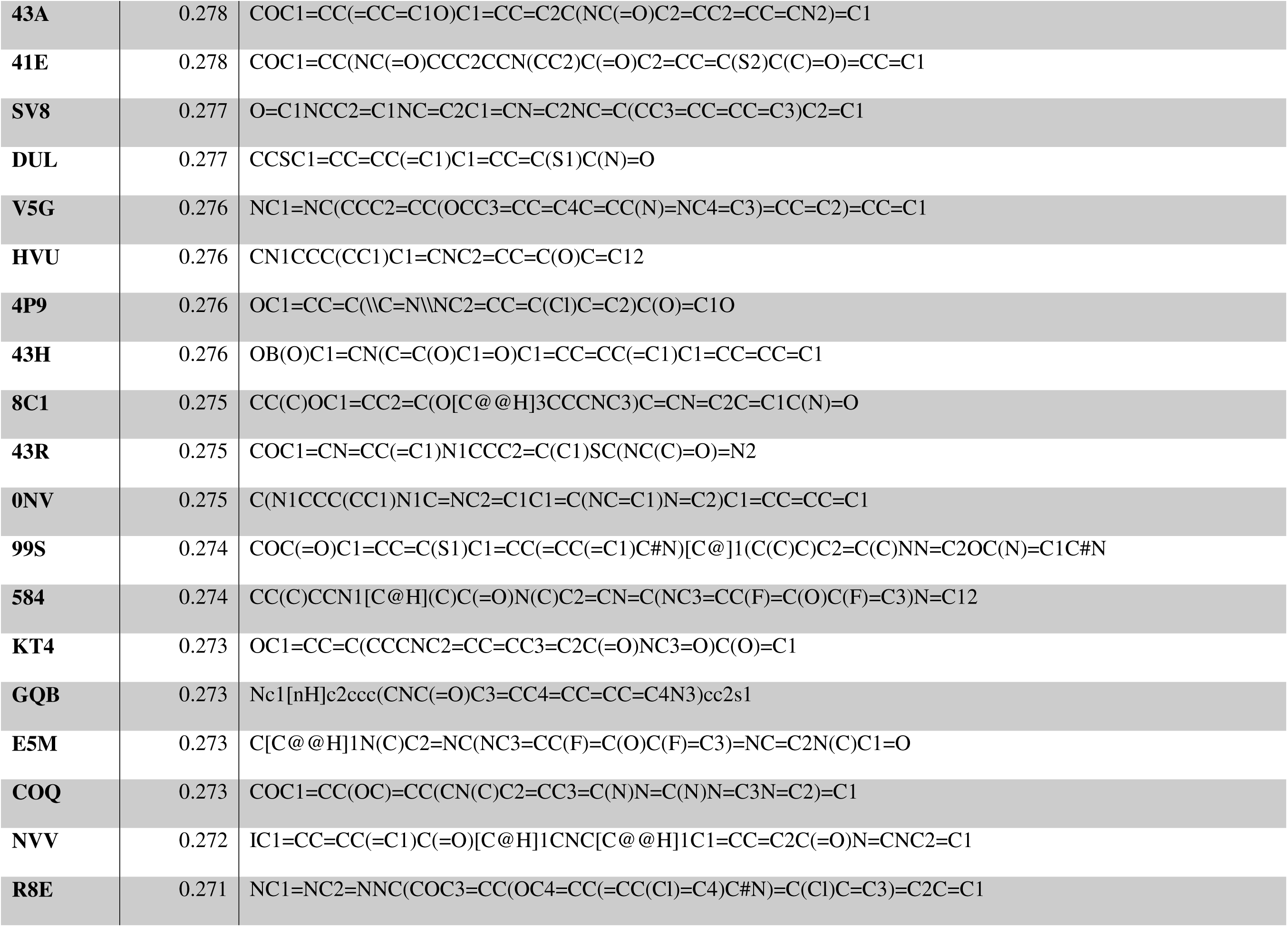
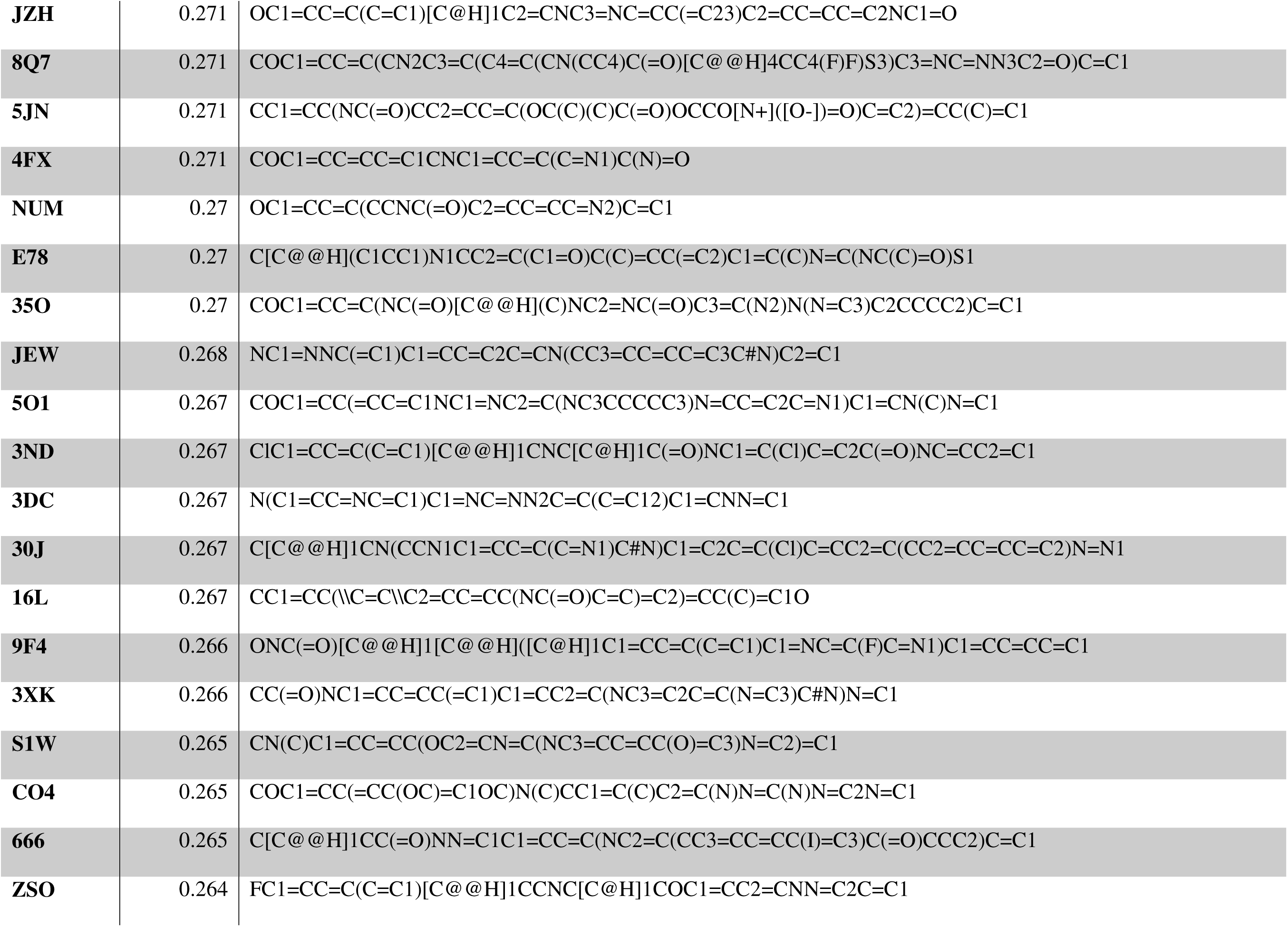
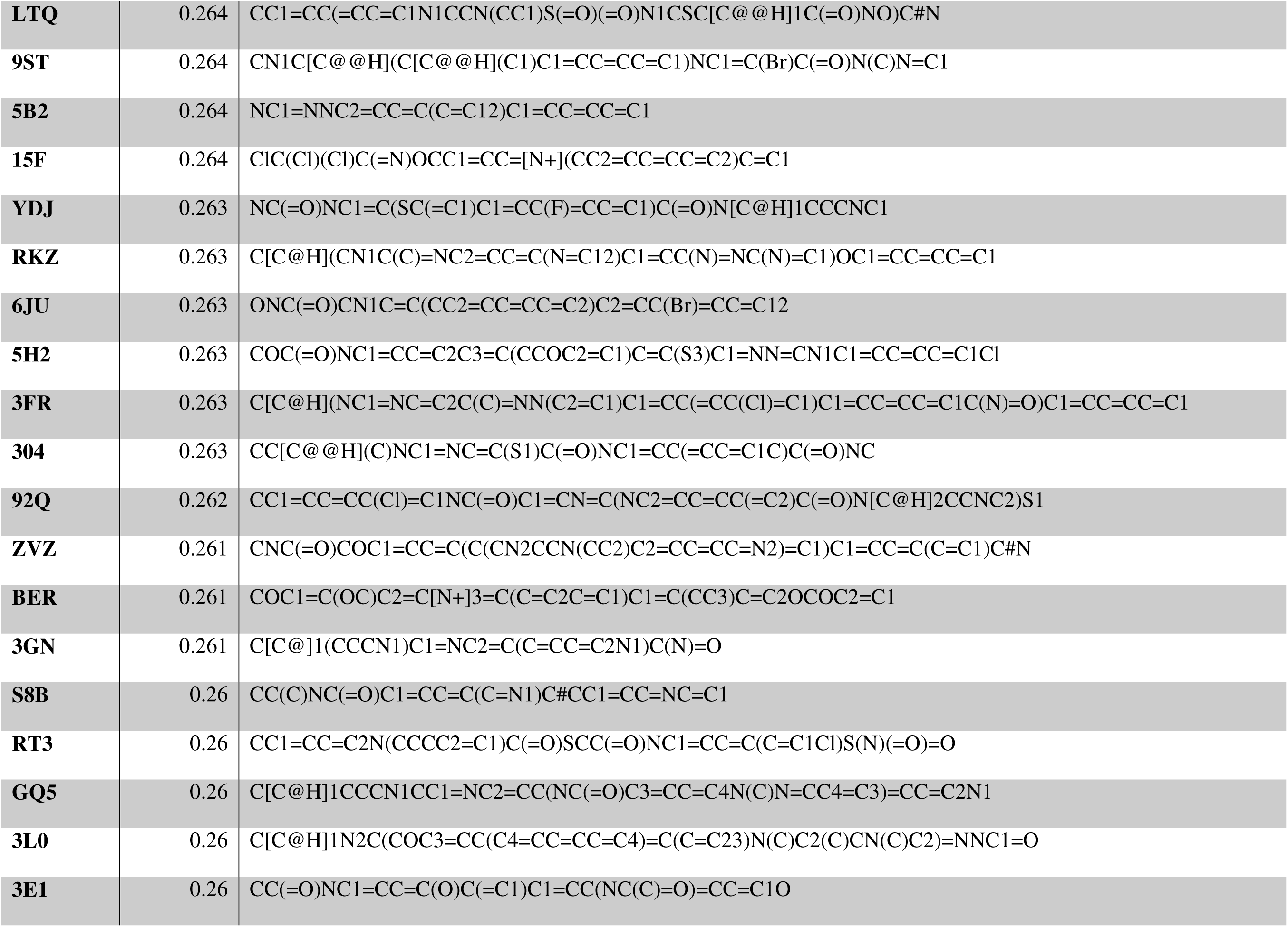
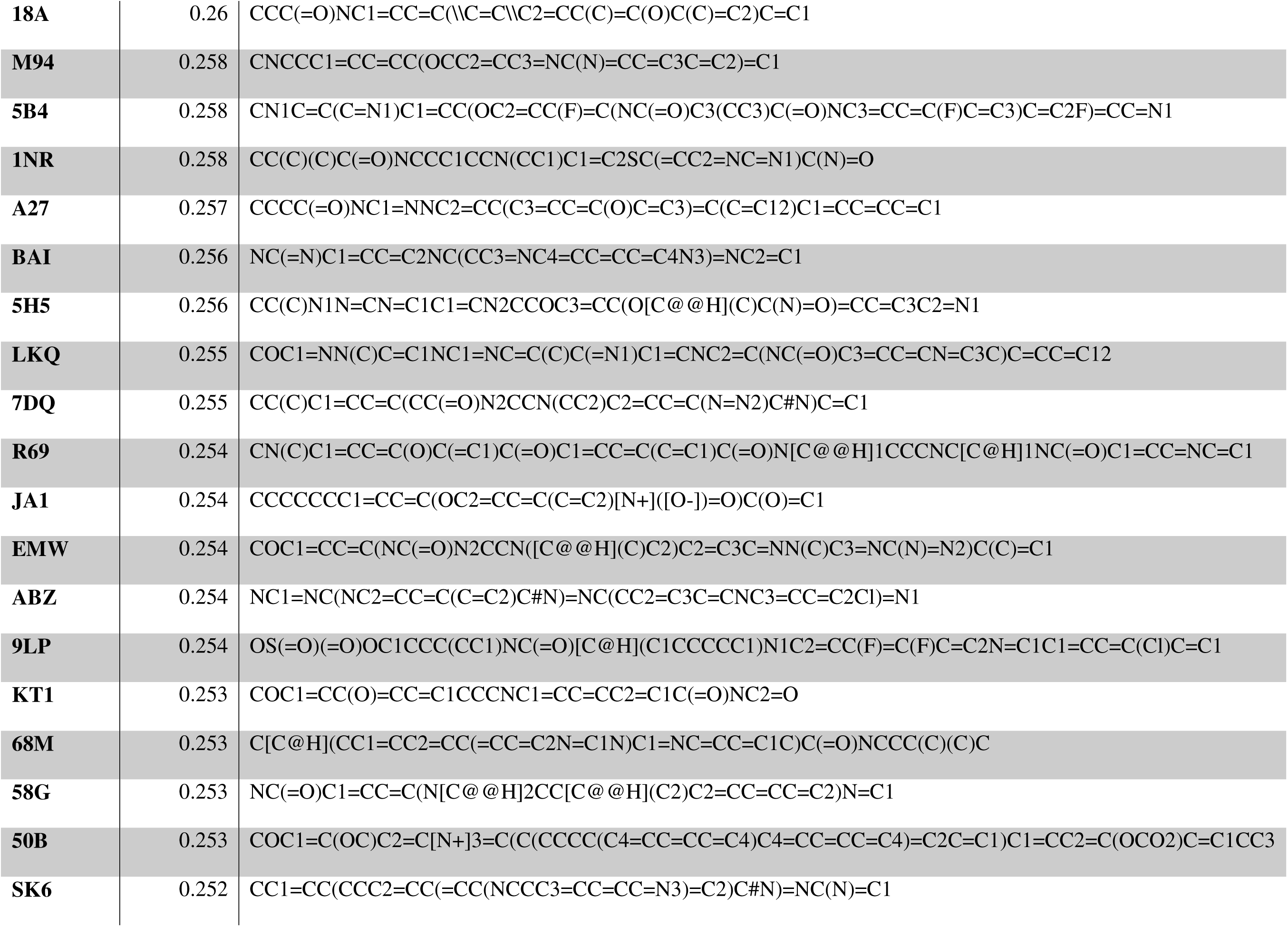
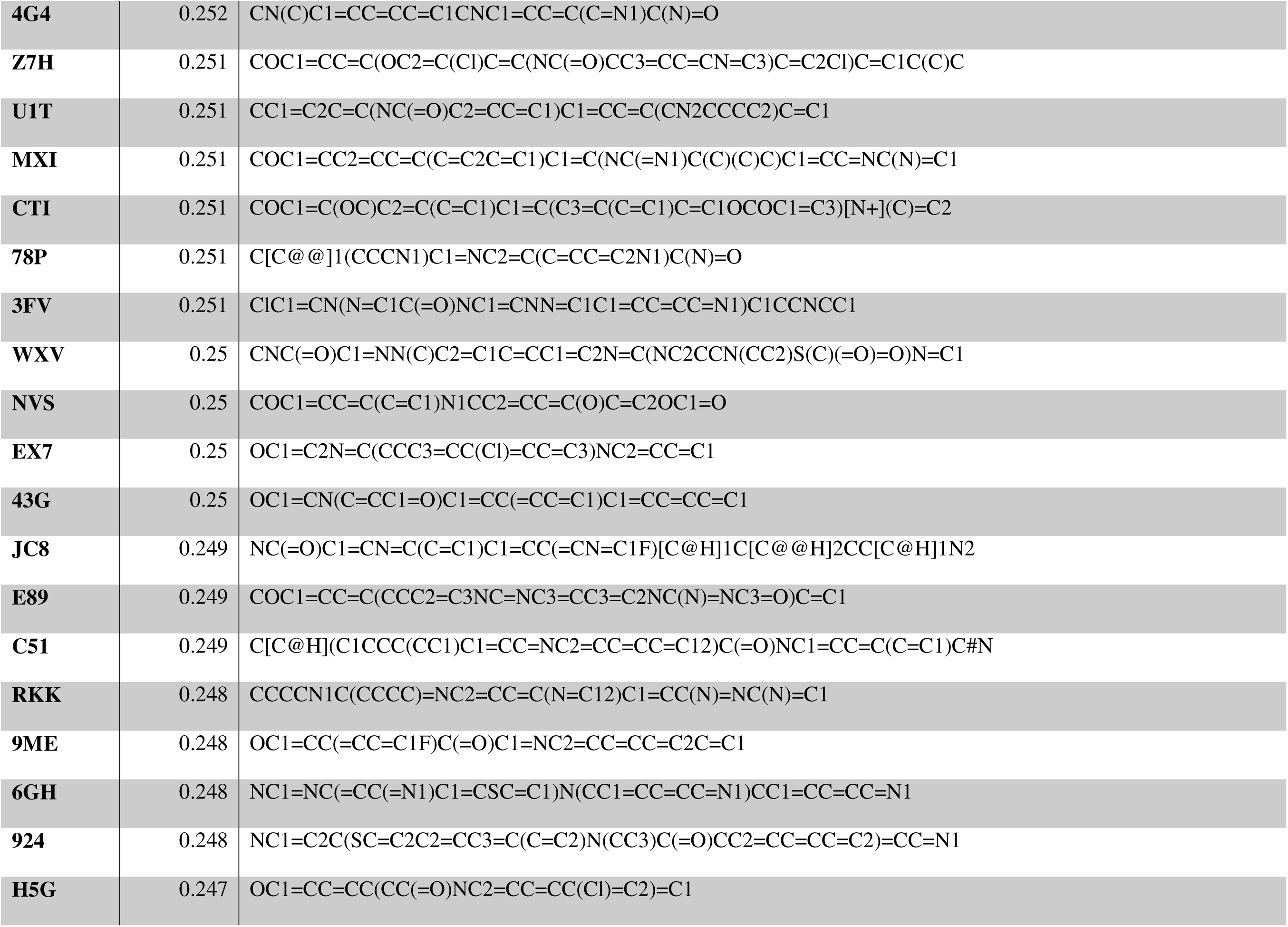
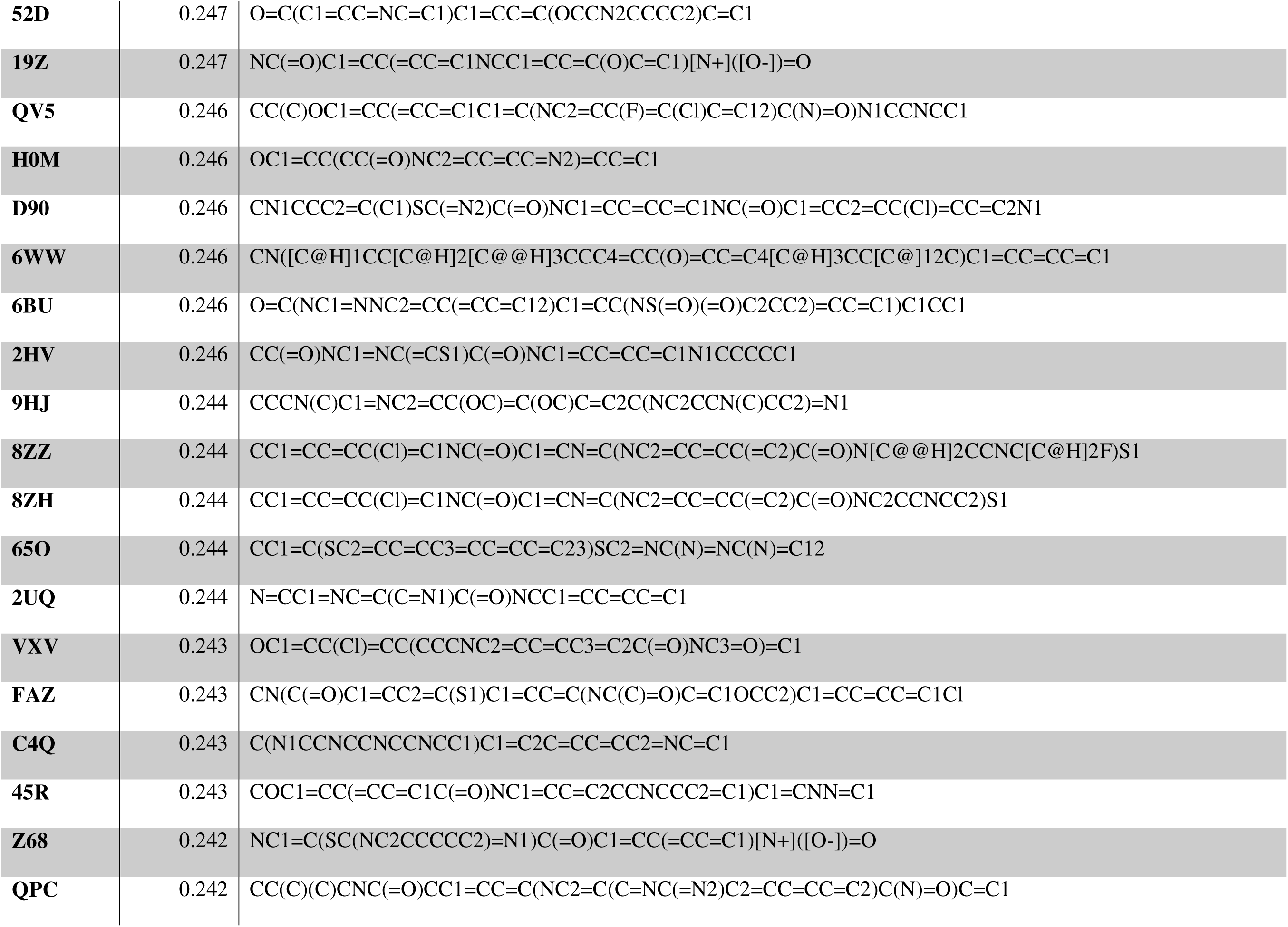
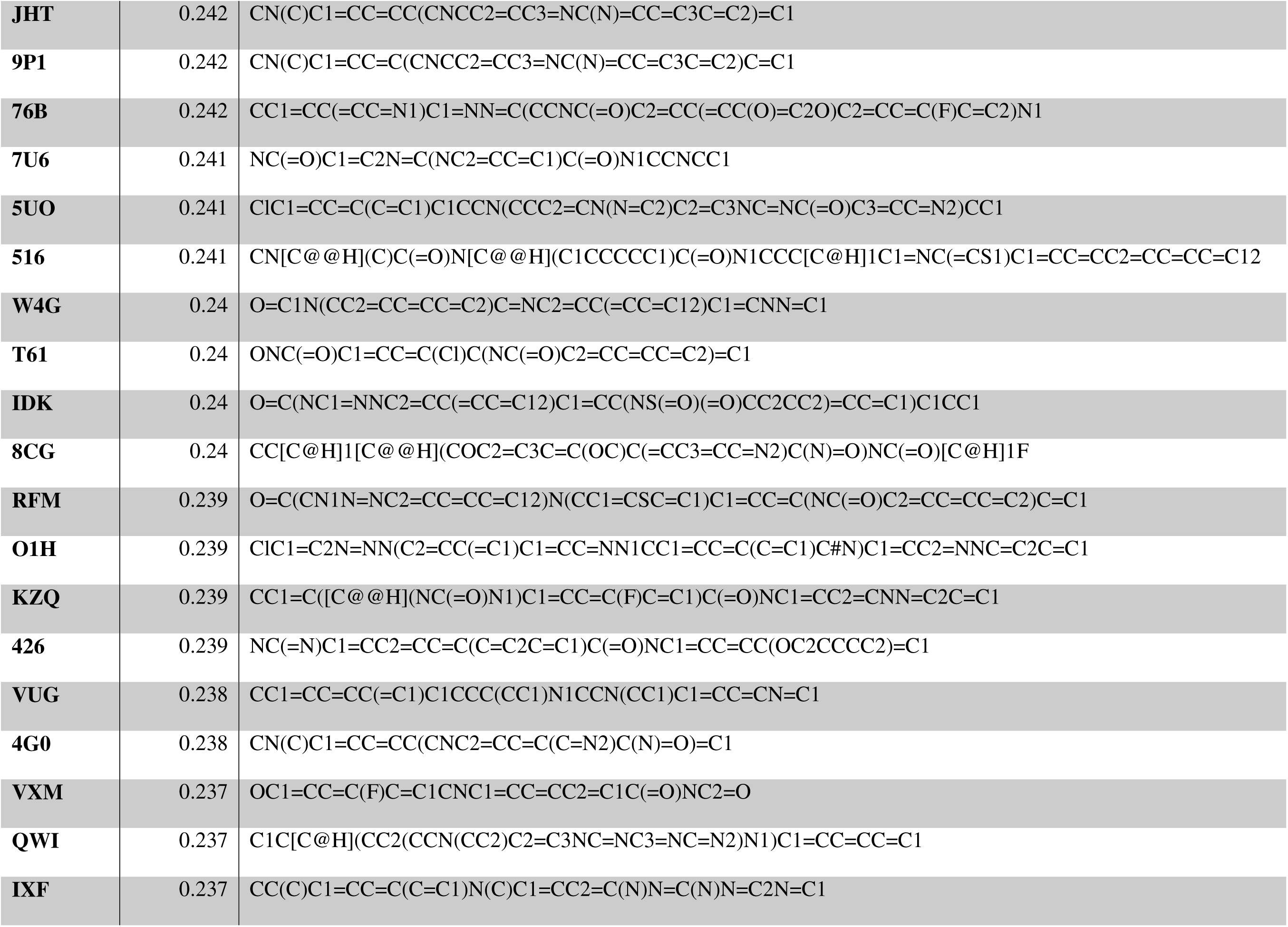
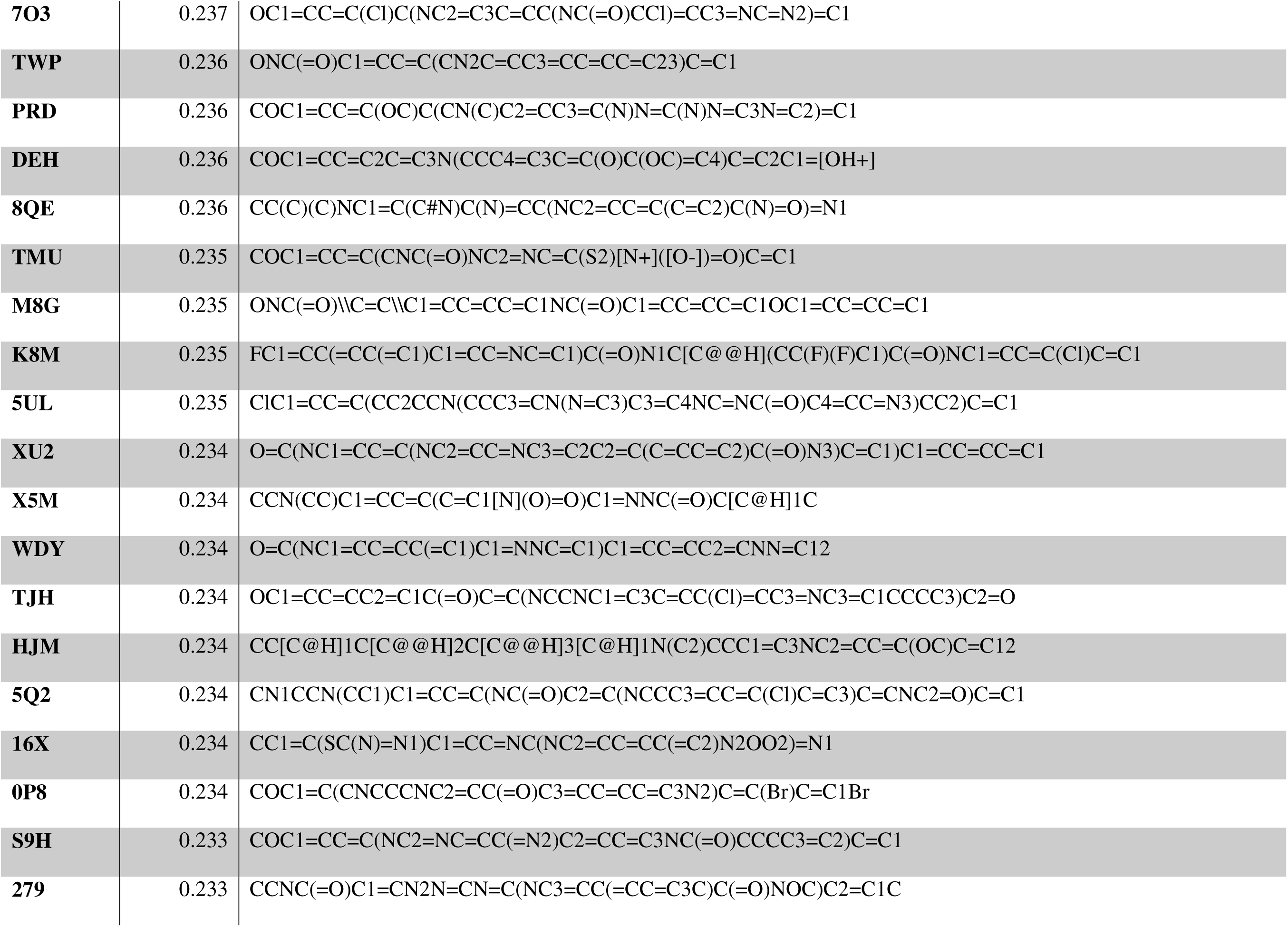
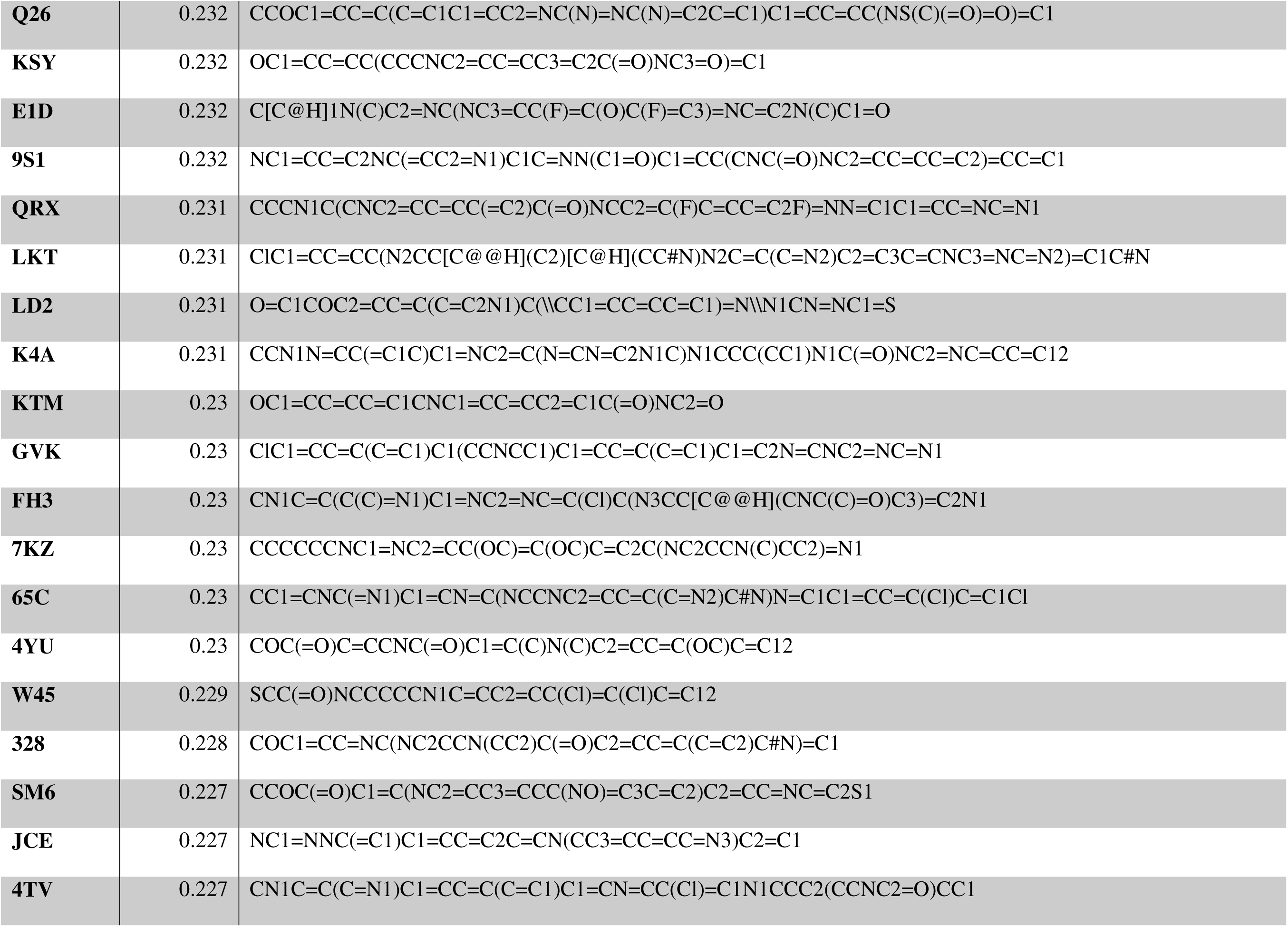
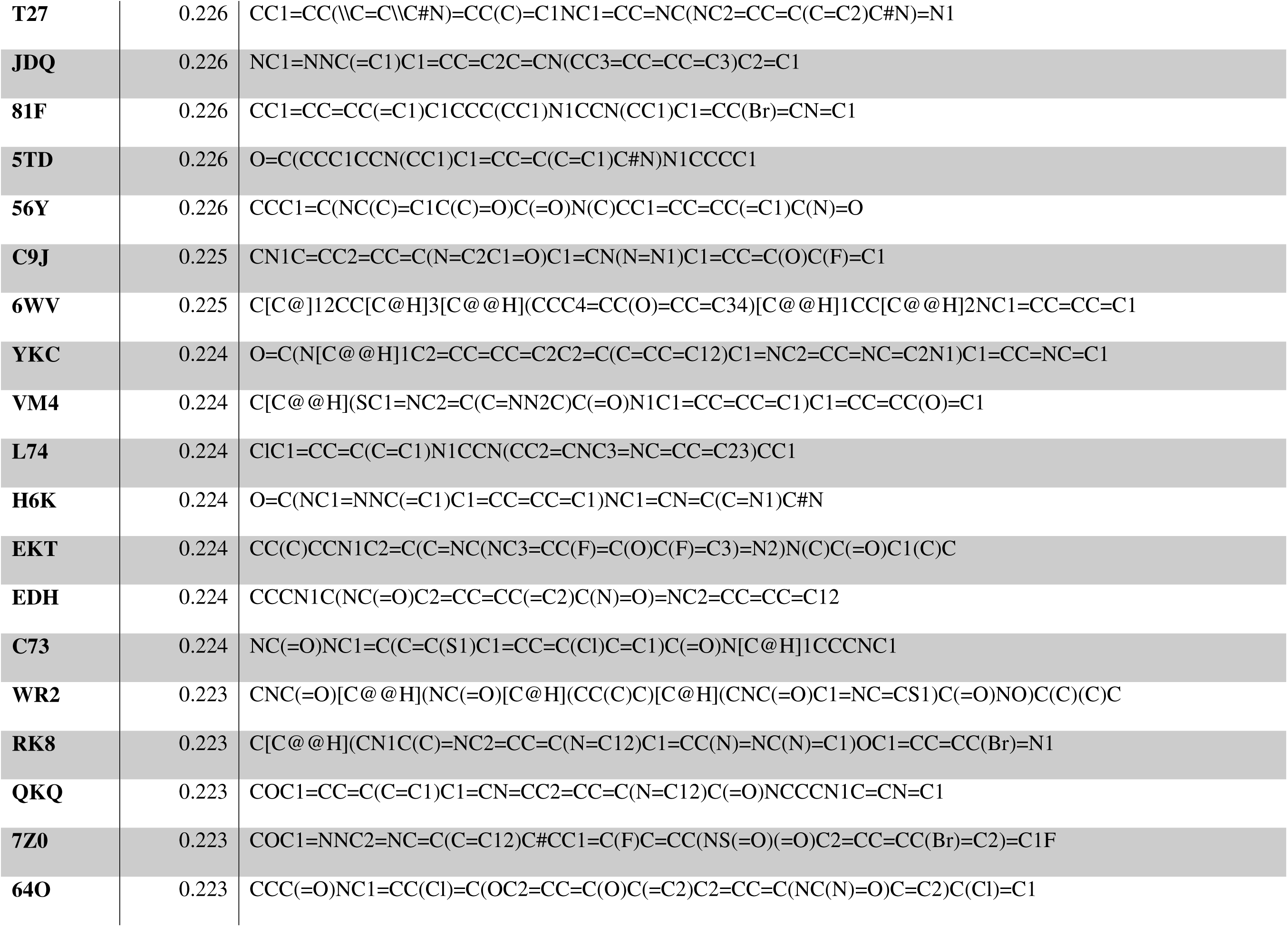
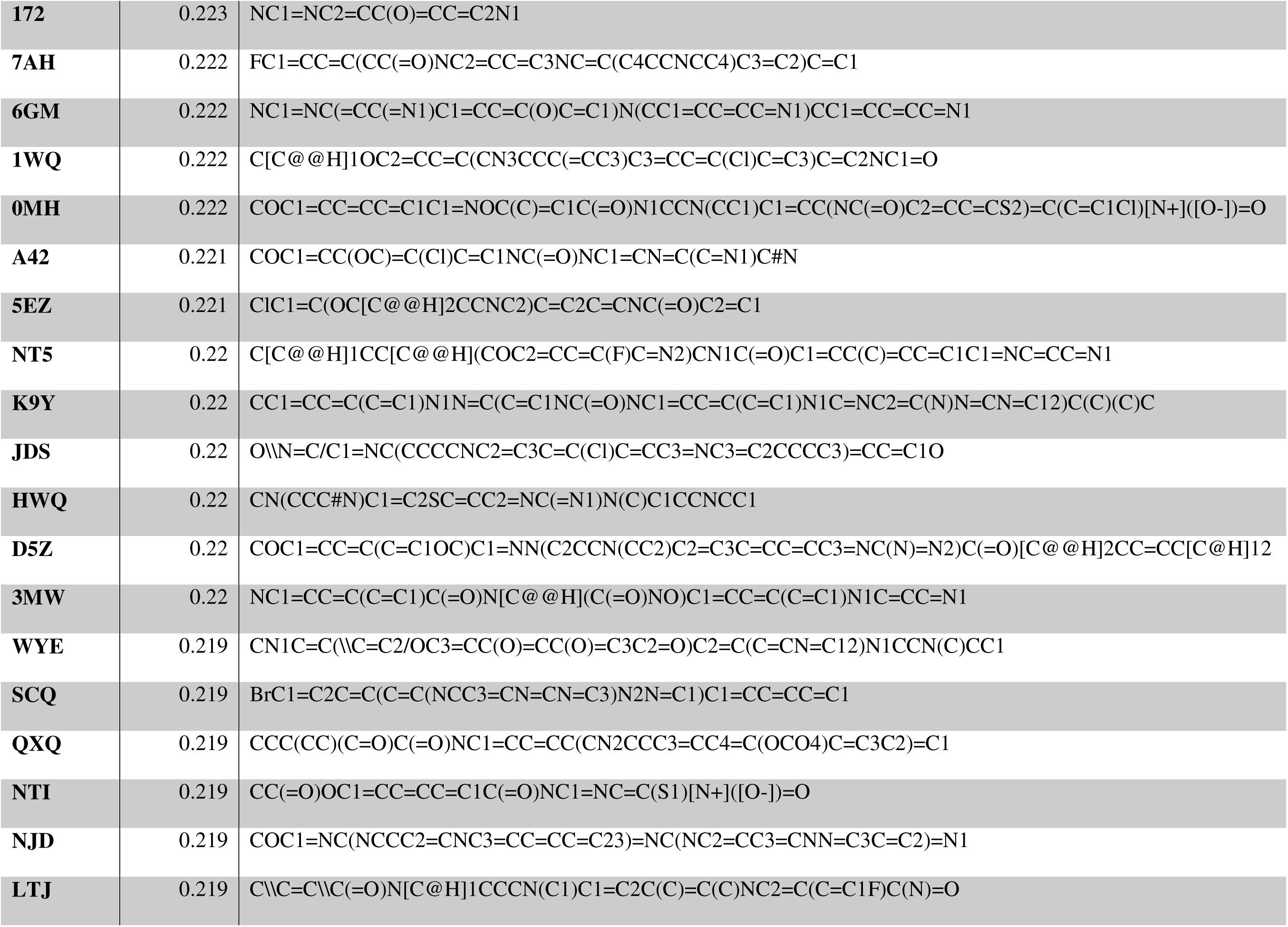
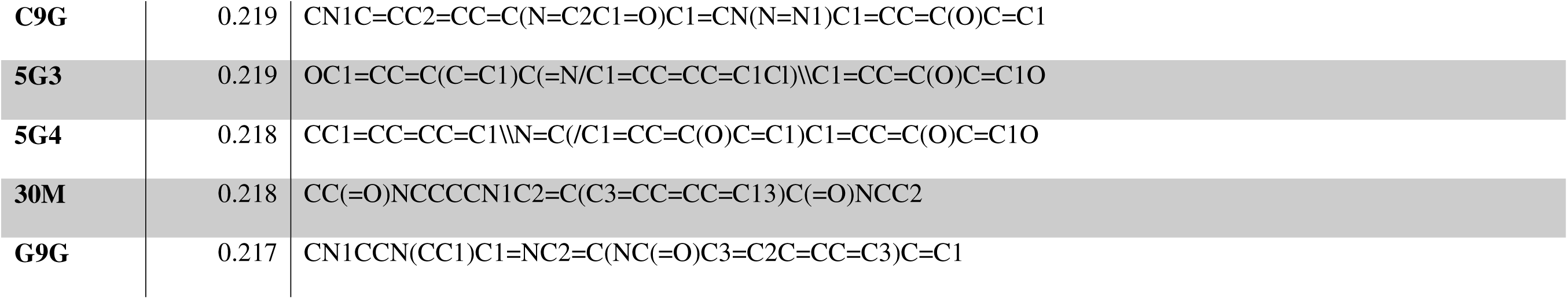

## References

(1) Nasrallah, I.; Dubroff, J. An Overview of PET Neuroimaging. Semin. Nucl. Med. 2013, 43 (6), 449–461. 10.1053/j.semnuclmed.2013.06.003.

(2) Burkett, B. J.; Babcock, J. C.; Lowe, V. J.; Graff-Radford, J.; Subramaniam, R. M.; Johnson, D. R. PET Imaging of Dementia: Update 2022. Clin. Nucl. Med. 2022, 47 (9), 763–773. 10.1097/RLU.0000000000004251.

(3) Wang, J.; Jin, C.; Zhou, J.; Zhou, R.; Tian, M.; Lee, H. J.; Zhang, H. PET Molecular Imaging for Pathophysiological Visualization in Alzheimer’s Disease. Eur. J. Nucl. Med. Mol. Imaging 2023, 50 (3), 765–783. 10.1007/s00259-022-05999-z.

(4) Jack, C. R.; Bennett, D. A.; Blennow, K.; Carrillo, M. C.; Feldman, H. H.; Frisoni, G. B.; Hampel, H.; Jagust, W. J.; Johnson, K. A.; Knopman, D. S.; Petersen, R. C.; Scheltens, P.; Sperling, R. A.; Dubois, B. A/T/N: An Unbiased Descriptive Classification Scheme for Alzheimer Disease Biomarkers. Neurology 2016, 87 (5), 539–547. 10.1212/WNL.0000000000002923.

(5) Rabinovici, G. D.; Knopman, D. S.; Arbizu, J.; Benzinger, T. L. S.; Donohoe, K. J.; Hansson, O.; Herscovitch, P.; Kuo, P. H.; Lingler, J. H.; Minoshima, S.; Murray, M. E.; Price, J. C.; Salloway, S. P.; Weber, C. J.; Carrillo, M. C.; Johnson, K. A. Updated Appropriate Use Criteria for Amyloid and Tau PET: A Report from the Alzheimer’s Association and Society for Nuclear Medicine and Molecular Imaging Workgroup. J. Nucl. Med. 2025. 10.2967/jnumed.124.268756.

(6) Aisen, P. S.; Cummings, J.; Jack, C. R.; Morris, J. C.; Sperling, R.; Frölich, L.; Jones, R. W.; Dowsett, S. A.; Matthews, B. R.; Raskin, J.; Scheltens, P.; Dubois, B. On the Path to 2025: Understanding the Alzheimer’s Disease Continuum. Alzheimers Res. Ther. 2017, 9 (1), 60. 10.1186/s13195-017-0283-5.

(7) Frisoni, G. B.; Altomare, D.; Thal, D. R.; Ribaldi, F.; van der Kant, R.; Ossenkoppele, R.; Blennow, K.; Cummings, J.; van Duijn, C.; Nilsson, P. M.; Dietrich, P.-Y.; Scheltens, P.; Dubois, B. The Probabilistic Model of Alzheimer Disease: The Amyloid Hypothesis Revised. Nat. Rev. Neurosci. 2022, 23 (1), 53–66. 10.1038/s41583-021-00533-w.

(8) Kepp, K. P.; Robakis, N. K.; Høilund-Carlsen, P. F.; Sensi, S. L.; Vissel, B. The Amyloid Cascade Hypothesis: An Updated Critical Review. Brain 2023, awad159. 10.1093/brain/awad159.

(9) Granzotto, A.; Sensi, S. L. Once upon a Time, the Amyloid Cascade Hypothesis. Ageing Res. Rev. 2024, 93, 102161. 10.1016/j.arr.2023.102161.

(10) Cole, G. B.; Keum, G.; Liu, J.; Small, G. W.; Satyamurthy, N.; Kepe, V.; Barrio, J. R. Specific Estrogen Sulfotransferase (SULT1E1) Substrates and Molecular Imaging Probe Candidates. Proc. Natl. Acad. Sci. U. S. A. 2010, 107 (14), 6222–6227. 10.1073/pnas.0914904107.

(11) Surmak, A. J.; Wong, K.-P.; Cole, G. B.; Hirata, K.; Aabedi, A. A.; Mirfendereski, O.; Mirfendereski, P.; Yu, A. S.; Huang, S.-C.; Ringman, J. M.; Liebeskind, D. S.; Barrio, J. R. Probing Estrogen Sulfotransferase-Mediated Inflammation with [11C]-PiB in the Living Human Brain. J. Alzheimers Dis. JAD 2020, 73 (3), 1023–1033. 10.3233/JAD-190559.

(12) Høilund-Carlsen, P. F.; Alavi, A.; Castellani, R. J.; Neve, R. L.; Perry, G.; Revheim, M.- E.; Barrio, J. R. Alzheimer’s Amyloid Hypothesis and Antibody Therapy: Melting Glaciers? Int. J. Mol. Sci. 2024, 25 (7), 3892. 10.3390/ijms25073892.

(13) Høilund-Carlsen, P. F.; Alavi, A.; Barrio, J. R. PET/CT/MRI in Clinical Trials of Alzheimer’s Disease. J. Alzheimer’s Dis. 2024, 101 (s1), S579–S601. 10.3233/JAD-240206.

(14) Cheung, J.; Rudolph, M. J.; Burshteyn, F.; Cassidy, M. S.; Gary, E. N.; Love, J.; Franklin, M. C.; Height, J. J. Structures of Human Acetylcholinesterase in Complex with Pharmacologically Important Ligands. J. Med. Chem. 2012, 55 (22), 10282–10286. 10.1021/jm300871x.

(15) Dvir, H.; Silman, I.; Harel, M.; Rosenberry, T. L.; Sussman, J. L. Acetylcholinesterase: From 3D Structure to Function. Chem. Biol. Interact. 2010, 187 (1–3), 10–22. 10.1016/j.cbi.2010.01.042.

(16) Auletta, J. T.; Johnson, J. L.; Rosenberry, T. L. Molecular Basis of Inhibition of Substrate Hydrolysis by a Ligand Bound to the Peripheral Site of Acetylcholinesterase. Chem. Biol. Interact. 2010, 187 (1–3), 135–141. 10.1016/j.cbi.2010.05.009.

(17) Harel, M.; Sonoda, L. K.; Silman, I.; Sussman, J. L.; Rosenberry, T. L. Crystal Structure of Thioflavin T Bound to the Peripheral Site of Torpedo Californica Acetylcholinesterase Reveals How Thioflavin T Acts as a Sensitive Fluorescent Reporter of Ligand Binding to the Acylation Site. J. Am. Chem. Soc. 2008, 130 (25), 7856–7861. 10.1021/ja7109822.

(18) Sulatskaya, A. I.; Rychkov, G. N.; Sulatsky, M. I.; Rodina, N. P.; Kuznetsova, I. M.; Turoverov, K. K. Thioflavin T Interaction with Acetylcholinesterase: New Evidence of 1:1 Binding Stoichiometry Obtained with Samples Prepared by Equilibrium Microdialysis. ACS Chem. Neurosci. 2018, 9 (7), 1793–1801. 10.1021/acschemneuro.8b00111.

(19) Alarcón-Enos, J.; Muñoz-Núñez, Evelyn; Gutiérrez, Margarita; Quiroz-Carreño, Soledad; Pastene-Navarrete, Edgar; and Céspedes Acuña, C. Dyhidro-β-Agarofurans Natural and Synthetic as Acetylcholinesterase and COX Inhibitors: Interaction with the Peripheral Anionic Site (AChE-PAS), and Anti-Inflammatory Potentials. J. Enzyme Inhib. Med. Chem. 2022, 37 (1), 1845–1856. 10.1080/14756366.2022.2091554.

(20) Singh, Y. P.; Shankar, G.; Jahan, S.; Singh, G.; Kumar, N.; Barik, A.; Upadhyay, P.; Singh, L.; Kamble, K.; Singh, G. K.; Tiwari, S.; Garg, P.; Gupta, S.; Modi, G. Further SAR Studies on Natural Template Based Neuroprotective Molecules for the Treatment of Alzheimer’s Disease. Bioorg. Med. Chem. 2021, 46, 116385. 10.1016/j.bmc.2021.116385.

(21) Sugimoto, H.; Yamanishi, Y.; Iimura, Y.; Kawakami, Y. Donepezil Hydrochloride (E2020) and Other Acetylcholinesterase Inhibitors. Curr. Med. Chem. 2000, 7 (3), 303–339. 10.2174/0929867003375191.

(22) David, B.; Schneider, P.; Schäfer, P.; Pietruszka, J.; Gohlke, H. Discovery of New Acetylcholinesterase Inhibitors for Alzheimer’s Disease: Virtual Screening and in Vitro Characterisation. J. Enzyme Inhib. Med. Chem. 2021, 36 (1), 491–496. 10.1080/14756366.2021.1876685.

(23) García-Ayllón, M.-S.; Small, D. H.; Avila, J.; Saez-Valero, J. Revisiting the Role of Acetylcholinesterase in Alzheimer’s Disease: Cross-Talk with P-Tau and β-Amyloid. Front. Mol. Neurosci. 2011, 4. 10.3389/fnmol.2011.00022.

(24) Campanari, M.-L.; Navarrete, F.; Ginsberg, S. D.; Manzanares, J.; Sáez-Valero, J.; García-Ayllón, M.-S. Increased Expression of Readthrough Acetylcholinesterase Variants in the Brains of Alzheimer’s Disease Patients. J. Alzheimer’s Dis. 2016, 53 (3), 831–841. 10.3233/JAD-160220.

(25) Glodzik, L.; Rusinek, H.; Li, J.; Zhou, C.; Tsui, W.; Mosconi, L.; Li, Y.; Osorio, R.; Williams, S.; Randall, C.; Spector, N.; McHugh, P.; Murray, J.; Pirraglia, E.; Vallabhajolusa, S.; de Leon, M. Reduced Retention of Pittsburgh Compound B in White Matter Lesions. Eur. J. Nucl. Med. Mol. Imaging 2015, 42 (1), 97–102. 10.1007/s00259-014-2897-1.

(26) Klunk, W. E.; Engler, H.; Nordberg, A.; Wang, Y.; Blomqvist, G.; Holt, D. P.; Bergstr??m, M.; Savitcheva, I.; Huang, G. F.; Estrada, S.; Aus??n, B.; Debnath, M. L.; Barletta, J.; Price, J. C.; Sandell, J.; Lopresti, B. J.; Wall, A.; Koivisto, P.; Antoni, G.; Mathis, C. A.; L??ngstr??m, B. Imaging Brain Amyloid in Alzheimer’s Disease with Pittsburgh Compound-B. Ann. Neurol. 2004, 55 (3), 306–319. 10.1002/ana.20009.

(27) Murugan, N. A.; Chiotis, K.; Rodriguez-Vieitez, E.; Lemoine, L.; Ågren, H.; Nordberg, A. Cross-Interaction of Tau PET Tracers with Monoamine Oxidase B: Evidence from in Silico Modelling and in Vivo Imaging. Eur. J. Nucl. Med. Mol. Imaging 2019, 46 (6), 1369–1382. 10.1007/s00259-019-04305-8.

(28) Zoete, V.; Daina, A.; Bovigny, C.; Michielin, O. SwissSimilarity: A Web Tool for Low to Ultra High Throughput Ligand-Based Virtual Screening. J. Chem. Inf. Model. 2016, 56 (8), 1399–1404. 10.1021/acs.jcim.6b00174.

(29) Bragina, M. E.; Daina, A.; Perez, M. A. S.; Michielin, O.; Zoete, V. The SwissSimilarity 2021 Web Tool: Novel Chemical Libraries and Additional Methods for an Enhanced Ligand-Based Virtual Screening Experience. Int. J. Mol. Sci. 2022, 23 (2), 811. 10.3390/ijms23020811.

(30) Kim, S.; Chen, J.; Cheng, T.; Gindulyte, A.; He, J.; He, S.; Li, Q.; Shoemaker, B. A.; Thiessen, P. A.; Yu, B.; Zaslavsky, L.; Zhang, J.; Bolton, E. E. PubChem 2023 Update. Nucleic Acids Res. 2023, 51 (D1), D1373–D1380. 10.1093/nar/gkac956.

(31) Floresta, G.; Granzotto, A.; Patamia, V.; Arillotta, D.; Papanti, G. D.; Guirguis, A.; Corkery, J. M.; Martinotti, G.; Sensi, S. L.; Schifano, F. Xylazine as an Emerging New Psychoactive Substance; Focuses on Both 5-HT7 and κ-Opioid Receptors’ Molecular Interactions and Isosteric Replacement. Arch. Pharm. (Weinheim) 2025, 358 (3), e2500041. 10.1002/ardp.202500041.

(32) Trott, O.; Olson, A. J. AutoDock Vina: Improving the Speed and Accuracy of Docking with a New Scoring Function, Efficient Optimization and Multithreading. J. Comput. Chem. 2010, 31 (2), 455. 10.1002/jcc.21334.

(33) Krieger, E.; Vriend, G. YASARA View - Molecular Graphics for All Devices - from Smartphones to Workstations. Bioinforma. Oxf. Engl. 2014, 30 (20), 2981–2982. 10.1093/bioinformatics/btu426.

(34) Krieger, E.; Dunbrack, R. L.; Hooft, R. W. W.; Krieger, B. Assignment of Protonation States in Proteins and Ligands: Combining pKa Prediction with Hydrogen Bonding Network Optimization. Methods Mol. Biol. Clifton NJ 2012, 819, 405–421. 10.1007/978-1-61779-465-0_25.

(35) Krieger, E.; Nielsen, J. E.; Spronk, C. A. E. M.; Vriend, G. Fast Empirical pKa Prediction by Ewald Summation. J. Mol. Graph. Model. 2006, 25 (4), 481–486. 10.1016/j.jmgm.2006.02.009.

(36) Maier, J. A.; Martinez, C.; Kasavajhala, K.; Wickstrom, L.; Hauser, K. E.; Simmerling, C. ff14SB: Improving the Accuracy of Protein Side Chain and Backbone Parameters from ff99SB. J. Chem. Theory Comput. 2015, 11 (8), 3696–3713. 10.1021/acs.jctc.5b00255.

(37) Wang, J.; Wolf, R. M.; Caldwell, J. W.; Kollman, P. A.; Case, D. A. Development and Testing of a General Amber Force Field. J. Comput. Chem. 2004, 25 (9), 1157–1174. 10.1002/jcc.20035.

(38) Jakalian, A.; Jack, D. B.; Bayly, C. I. Fast, Efficient Generation of High-Quality Atomic Charges. AM1-BCC Model: II. Parameterization and Validation. J. Comput. Chem. 2002, 23 (16), 1623–1641. 10.1002/jcc.10128.

(39) Hornak, V.; Abel, R.; Okur, A.; Strockbine, B.; Roitberg, A.; Simmerling, C. Comparison of Multiple Amber Force Fields and Development of Improved Protein Backbone Parameters. Proteins 2006, 65 (3), 712–725. 10.1002/prot.21123.

(40) Essmann, U.; Perera, L.; Berkowitz, M. L.; Darden, T.; Lee, H.; Pedersen, L. G. A Smooth Particle Mesh Ewald Method. J. Chem. Phys. 1995, 103 (19), 8577–8593. 10.1063/1.470117.

(41) Krieger, E.; Vriend, G. New Ways to Boost Molecular Dynamics Simulations. J. Comput. Chem. 2015, 36 (13), 996–1007. 10.1002/jcc.23899.

(42) Floresta, G.; Patamia, V.; Mazzeo, P. P.; Lombardo, G. M.; Pistarà, V.; Bacchi, A.; Rescifina, A.; Punzo, F. Structural, Morphological, and Modeling Studies of N- (Benzoyloxy)Benzamide as a Specific Inhibitor of Type II Inosine Monophosphate Dehydrogenase. J. Mol. Struct. 2024, 1303, 137588. 10.1016/j.molstruc.2024.137588.

(43) Granzotto, A.; Bolognin, S.; Scancar, J.; Milacic, R.; Zatta, P. Beta-Amyloid Toxicity Increases with Hydrophobicity in the Presence of Metal Ions. Monatshefte Chem. 2011, 142 (4), 421–430. 10.1007/s00706-011-0470-1.

